# Enhancement of activation-induced T cell proliferation by SIRPG in a CD47-independent manner

**DOI:** 10.1101/2025.05.01.651731

**Authors:** Flavien Marguerie, Mohammad A. Saifi, Benjamin Geary, David Barnes, Anna H. Jonsson, I-Cheng Ho

**Affiliations:** Division of Rheumatology, Inflammation, and Immunity, Department of Medicine, Brigham and Women’s Hospital, Boston, MA 02115; Harvard Medical School, Boston, MA 02115

## Abstract

SIRPG, a primate-specific type 1 transmembrane protein in the Signal Regulatory Protein (SIRP) family, is predominantly expressed in T cells. It contains a short cytoplasmic domain, which does not contain any known signaling motif, and its only known ligand is CD47. Several genetic variations in *SIRPG*, including the V263A (rs6043409) polymorphism, linked to increased type 1 diabetes risk, highlight its potential importance. However, its expression and physiological role remain largely unclear due to its absence in rodents. Here, we demonstrate that SIRPG and GzmB exhibit near mutually exclusive expression in resting peripheral CD8+ T cells. We further show that SIRPG serves as a valuable marker for GzmK-expressing CD8+ T cells in peripheral blood and inflamed synovial fluid and that its expression in both CD4+ and CD8+ T cells is upregulated by anti-CD3 stimulation, with further enhancement by the TNFα inhibitor adalimumab, but not certolizumab. While SIRPG ablation minimally affects T cell activation and IFNγ/TNFα production, it impairs the expression of mitosis-regulating genes like *UBE2C* and *TOP2A*, leading to reduced proliferation, and alters the expression of certain activation-induced surface molecules, including CRTAM. Notably, SIRPG-mediated proliferation and CRTAM expression are cell-autonomous and CD47-independent. Structural and functional analyses reveal that SIRPG-driven proliferation is independent of its extracellular D1 domain, not significantly affected by the V263 variant, but dependent on its cytoplasmic domain. Collectively, our findings offer novel insights into the expression, function, and mechanism of action of SIRPG in T cells.

## Introduction

SIRPG is a member of the Signal Regulatory Protein (SIRP) family of type 1 transmembrane proteins, which in humans also includes SIRPA, SIRPB1, SIRPB2, and SIRPD (1, 2). These SIRPs are characterized by three extracellular immunoglobulin-like domains (D1-3, N-terminus to C-terminus), a transmembrane domain, and a cytoplasmic domain of varying length. Notably, only SIRPA has a murine homolog. Predominantly expressed in myeloid cells, SIRPA possesses both ITIM and ITSM motifs in its cytoplasmic domain. Its well-characterized ligand, CD47, is ubiquitously expressed, and their interaction delivers a “don’t-eat-me” signal in myeloid cells. Cancer cells often exploit this inhibitory SIRPA-CD47 axis to evade phagocytosis. Murine SIRPB and human SIRPB1/2 are also primarily found on myeloid cells. Their short intracellular domains lack known signaling motifs, but their transmembrane domains contain a conserved lysine residue. Research on murine SIRPB has shown that this lysine is crucial for association with DAP12 (3), which contains a single ITAM motif, suggesting a potential activating role for SIRPBs through DAP12. Indeed, cross-linking murine SIRPB triggers Syk and MAPK phosphorylation in myeloid cells (4).

In contrast to other SIRP family members, SIRPG exhibits near-exclusive expression in primate T and NK cells (5). It is highly expressed in naive (Tnaive) and central memory (Tcm) T cells, with expression further upregulated upon anti-CD3 stimulation but downregulated in effector memory (Tem) and CD45RA+ effector memory (Temra) T cells. This evolutionary late appearance and restricted expression suggest a specialized role for SIRPG in primate adaptive immunity. Despite this, a comprehensive understanding of its expression and function is still lacking. Several single nucleotide polymorphisms (SNPs) within the *SIRPG* locus are associated with increased susceptibility to type 1 diabetes (T1D) and narcolepsy (6–10). Notably, a missense SNP (rs6043409) encoding a V263A variation in the D3 domain is predicted to induce conformational changes and is likely causal (7). Furthermore, elevated *SIRPG* transcript levels in PBMCs and/or protein levels in serum have been observed in patients with T1D, diabetic retinopathy, and active lupus (but not inactive lupus) (11–16), suggesting SIRPG promotes immune responses and contributes to autoimmune pathogenesis. Conversely, the risk allele of another T1D-associated non-coding SNP (rs2281808) correlates with reduced *SIRPG* transcript levels in the thymus and surface protein levels on peripheral T cells (17, 18). Additionally, a higher percentage of SIRPG^low^ CD8+ peripheral T cells has been reported in patients with T1D and relapsing-remitting multiple sclerosis (RRMS), independent of genotype (19), implying SIRPG may inhibit adaptive immunity and offer protection in autoimmune diseases. The underlying reasons for these conflicting findings remain unclear.

Published in vitro investigations into SIRPG function have produced inconclusive and sometimes contradictory findings for several key reasons. First, while CD47 is the only identified ligand for SIRPG, it also binds to other proteins, including SIRPA, integrins, and TSP-1, complicating the interpretation of CD47-based assays. Second, the relatively lower affinity of CD47 for SIRPG compared to SIRPA suggests the potential existence of additional, yet undiscovered, SIRPG ligands. Third, the reagents frequently employed in these studies, such as anti-SIRPG antibodies, anti-CD47 antibodies, and CD47-Fc fusion proteins, may lack full validation and likely exhibit off-target effects, contributing to inconsistent results. Furthermore, SIRPG’s short cytoplasmic domain (4 amino acids) and lack of a DAP12-interacting lysine in its transmembrane domain have led some to hypothesize that it functions solely as an adhesion molecule without signaling capacity. However, evidence suggests otherwise: anti-CD47 or anti-SIRPG (LSB2.20) partially inhibits T cell proliferation and IFNγ production induced by APC/superantigen or allogeneic immature DCs (5), presumably by disrupting SIRPG-CD47 interactions. Conversely, soluble LSB2.20 or plate-bound CD47-Fc enhances the viability/proliferation and IFNγ production of anti-CD3-activated T cells (5, 20), potentially through SIRPG engagement. Additionally, SIRPG recruitment to the immune synapse between Raji B cells and Jurkat T cells in the presence of a superantigen suggests its involvement in signaling (20). Nevertheless, discrepancies persist. For instance, the co-stimulatory effect of CD47-Fc on T cells is only partially blocked by SIRPG inhibition (20), indicating SIRPG-independent mechanisms. Moreover, engaging CD47 on Jurkat T cells with SIRPG-coated beads induces apoptosis (21), suggesting that SIRPG-CD47 interactions among T cells can trigger opposing bidirectional signals: activation via SIRPG and apoptosis via CD47. How T cells integrate these conflicting signals remains an open question.

In vivo studies have also yielded inconsistent results. Administration of a pan-SIRP antibody (KWAR23) reduces the engraftment of human T cells in NSG (NOD-SCID-gamma) mice, a xenogenic graft-versus-host (xenoGVHD) model, and enhances the survival of animals (20). However, because KWAR23 can block the interaction between SIRPA and CD47, its in vivo effect cannot be definitely attributed to SIRPG inhibition. The at-risk T allele of the C-to-T SNP rs2281808 is associated with higher percentage of SIRPG^low^ cells in PBMC (18, 22). These SIRPG^low^ cells from healthy CT donors exhibit an effector phenotype, lower activation threshold in response to anti-CD3 in vitro, and greater pathogenicity in inducing xenoGVHD in NSG mice compared to SIRPG^high^ T cells (19). This seemingly contradicts the KWAR23 findings. However, the observed differences between SIRPG^high^ and SIRPG^low^ cells might be attributable to the SIRPG^low^ population containing a higher proportion of Tem and Temra cells, which naturally express lower SIRPG levels, compared to the SIRPG^high^ population enriched in naive and Tcm cells.

Given the caveats of the existing SIRPG-targeting reagents and the intrinsic differences between SIRPG^low^ and SIRPG^high^ cells, we opted to use CRISPR to ablate SIRPG in peripheral T cells of healthy donors and performed unbiased bulk RNA-seq analysis to investigate the function of SIRPG. Structural and functional analyses were subsequently carried out to determine the extent to which SIRPG’s activity depends on its extracellular and cytoplasmic domains.

## Results

### Expression of SIRPG in resting PBMC

It has been reported that Temra and Tem cells expressed a lower level of SIRPG. To further define the expression of SIRPG in various subsets of peripheral T cells, we used CD45RO, CD27, CD45RA, and CD62L to define the phenotype of resting peripheral CD4+ T cells from healthy donors (Figure S1A), and then examined the expression of SIRPG, CD28, and CCR7 in each subset. We found that the “naïve” subset (CD45RO^−^CD27^+^) already expressed a high level of SIRPG regardless of the status of CD45RA or CD62L (Figure 1A). The level was slightly increased in “Tcm” subset (CD45RO^+^CD27^+^). The transition from SIRPG^hi^ to SIRPG^−^ started in “Tem” cells (CD45RO^+^CD27^−^); a significant fraction of CD45RO^+^CD27^−^CD45RA^−^CD62L^−^ Tem cells were SIRPG^low^ or SIRPG^−^. The level continued to drop through the CD45RO^−^CD27^−^CD45RA^−^CD62L^−^ subset, whereas the classical Temra (CD45RO^−^CD27^−^CD45RA^+^) became SIRPG^−^, a finding in agreement with the published data. This continuum of expression of SIRPG correlated strongly with that of CD28 but was in contrast to that of CCR7, whose level in Temra was very close to that of naïve T cells. A similar pattern of expression of SIRPG was observed in resting peripheral CD8+T cells (Figure S1B and S1C).

**Figure 1.**
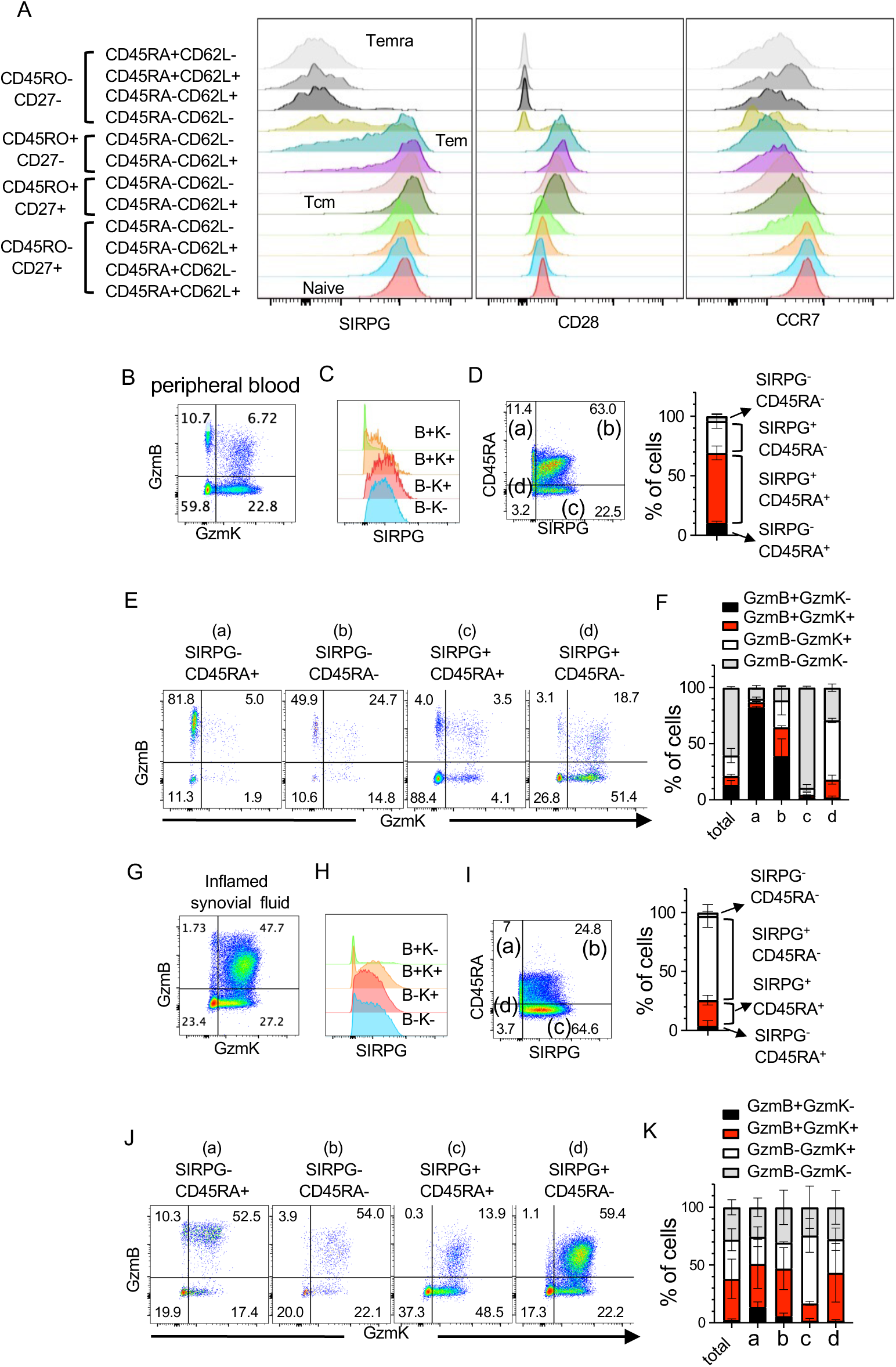
Expression of SIRPG in peripheral blood mononuclear cells and synovial fluid cells. **A**. PBMC from healthy donors were stained with antibodies against indicated surface markers and analyzed with FACS. The levels of SIRPG, CD28, and CCR7 of indicated populations are shown in overlay histograms in **A**. **B-K**. PBMC from two healthy donors (**B-F**) and synovial fluid cells from two patients with inflammatory arthritis (**G-K**) were stained for surface SIRPG and CD45RA and intracellular GzmK and GzmB. The stained cells were first separated into four populations based on the expression of GzmK and GzmB (**B** & **G**). The levels of SIRPG in the four GzmK/GzmB populations are shown in overlay histograms (**C** & **H**). The stained PBMC and synovial fluid cells were also separated into four populations based on the expression of SIRPG and CD45RA (**D** & **I**). The percentages of the SIRPG/CD45RA populations are shown in the cumulated bar graphs (**D** & **I**). The expression of GzmK and GzmB in each SIRPG/CD45RA population is then shown in the dot plots in **E** & **J**. The percentage of cells positive or negative for GzmB or GzmK from the SIRPG/CD45RA populations is shown in the cumulative bar graph of **F** and **K**.

### Identifying Granzyme B^+^ and Granzyme K^+^ CD8+ T cells by surface expression of SIRPG

Granzyme K (GzmK)-expressing CD8+T cells were recently found to be enriched in several autoimmune and inflammatory diseases (23), including rheumatoid arthritis. They are functionally distinct from GzmB-expressing cells and play a pathogenic role in autoimmune and inflammatory diseases. Thus far, there is no reliable surface marker to distinguish between GzmK^+^ and GzmB^+^ T cells. A recent RNA-seq analysis of peripheral T cells indicates that GzmB was highly expressed in SIRPG^low^ peripheral CD8+T cells compared to SIRPG^hi^ CD8+T cells (18). We therefore examined the expression of surface SIRPG, intracellular GzmK and GzmB in PBMC from healthy donors. Approximately 11%, 7%, 20%, and 60% of CD8+T cells were GzmB^+^GzmK^−^, GzmB^+^GzmK^+^, GzmB^−^GzmK^+^, and GzmB^−^GzmK^−^ cells, respectively (Figure 1B). Interestingly, the great majority of GzmB^−^GzmK^−^ and GzmB^+^GzmK^−^ cells were SIRPG^+^ and SIRPG^−^, respectively (Figure 1C). As naive cells, which are typically CD45RA^+^GzmB^−^GzmK^−^, we therefore used SIRPG and CD45RA to divide peripheral CD8+T cells into four populations: SIRPG^−^CD45RA^+^ (12%), SIRPG^−^CD45RA^−^ (3%), SIRPG^+^CD45RA^+^ (60%), and SIRPG^+^CD45RA^−^ (22%) (Figure 1D). More than 80% of the SIRPG^−^CD45RA^+^ cells were GzmB single positive cells; approximately 90% of the SIRPG^+^CD45RA^+^ cells expressed neither GzmK nor GzmB; and nearly 70% of the SIRPG^+^CD45RA^−^ cells expressed GzmK (only or together with GzmB) (Figure 1E and 1F). Thus, the expression of SIRPG and GzmB is almost mutually exclusive and the combination of SIRPG and CD45RA facilitates the distinction among GzmB^+^GzmK^−^ (enriched in population a), GzmB^−^ GzmK^−^ (enriched in population c), and GzmK^+^ (enriched in population d) CD8+T cells of peripheral blood. However, this distinction was blurred in inflamed synovial fluid. There were much fewer GzmB^+^GzmK^−^ cells (1%-3%) (Figure 1G), the greater majority of which were still SIRPG^−^ (Figure 1H), and a marked increase in the percentage of GzmB^+^GzmK^+^ cells (23%-47%) and downregulation of CD45RA (Figure 1G and 1I). In addition, there was wider variation between two donors, making SIRPG less reliable in distinguishing between GzmB^+^ and GzmK^+^ cells. Nevertheless, the SIRPG^+^CD45RA^+^ gate (population c) still partly excluded GzmB-expressing cells, thereby modestly enriching GzmB^−^GzmK^+^ cells (Figure 1J and 1K).

### Differential effects of TNFα inhibitors on the expression of SIRPG

Previous studies have shown that the level of surface SIRPG in bulk Th cells increased after they underwent two cycles of division (20) and that its expression was enhanced by Fc-containing TNFα inhibitors adalimumab (ada) and etanercept (eta) (24), but not certolizumab (cer), which lacks Fc. In agreement with these observations, we found that the surface level of SIRPG in CD4+ and CD8+ T cells of anti-CD3 stimulated PMBC increased over a course of 6 day and was further induced by anti-CD3 re-stimulation (Figure 2A and Figure S2A). The effect of ada when added at time 0 became obvious 72 hours after stimulation when examined with FACS, was detected in both blasting (high FSC) (Figure S2B and 2B) and non-blasting (low FSC) (Figure S2B and 2C) CD4+ but less obvious in non-blasting CD8+ T cells of PBMC (Figure 2B and 2C). Eta, but not cer or tocilizumab (toc, a humanized IgG1 against human IL-6R), also showed a strong trend of enhancing the expression of SIRPG in both CD4 and CD8 T cells (Figure 2B and 2C). In addition, resting T cells once encountered ada during the initial stimulation with anti-CD3/anti-CD28 maintained a higher level of SIRPG up to day 6 and upon re-stimulation with anti-CD3 compared to those without encountering ada during the initial stimulation (Figure 2D). Ada still enhanced, albeit to a less degree, the expression of SIRPG when added 72 hours after the initial ant-CD3 stimulation (Figure 2E). This lesser effect was probably due to a high level of SIRPG at this time point even before adding ada. Interestingly, ada did not augment the expression of SIRPG in resting T cells from un-stimulated PBMC (Figure 2F). As resting peripheral T cells included antigen-experienced SIRPG^−^ Temra and/or Tem, the effect of ada on the expression of SIRPG is dependent on T cell activation regardless of the history of antigen exposure. By contrast, none of the drugs altered the level of CD4, CD8, or the CD4/CD8 ratio (Figure S2C)

**Figure 2.**
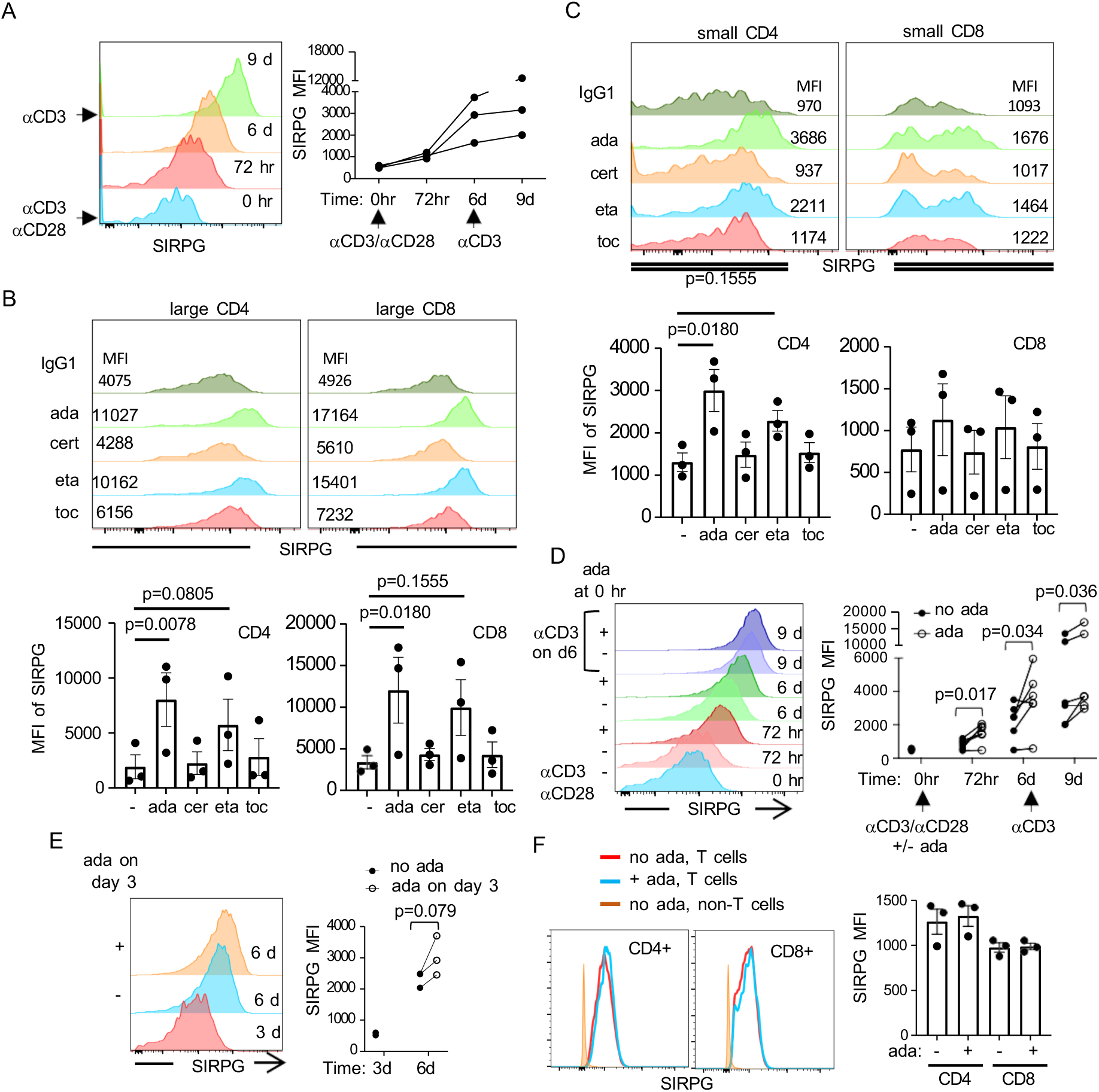
Induction of SIRPG by adalimumab but not certolizumab. **A**. PBMC from healthy donors were stimulated with anti-CD3/anti-CD28 and restimulated with anti-CD3 on day 6. The surface level of SIRPG in CD4+ T cells was analyzed with FACS and shown in the overlay histograms (the left panel). The MFI of SIRPG from three independent donors over the time course is shown in the right panel. Adalimumab (ada), certolizumab (cer), etanercept (eta), tocilizumab (toc), or control IgG1 was added on day 0 during the stimulation. The MFI of SIRPG from non-blasting (**B**) and blasting (**C**) T cells is shown in the overlay histograms. The cumulative results from 3 independent experiments are shown in the corresponding bar graphs. **D**. PBMC were stimulated in the presence or absence of ada and the surface level of SIRPG was monitored with FACS at indicated time points. Representative data from one donor is shown in the overlay histograms and cumulated results from five donors are shown in the right panel. The data points from the same donors are connected with lines. **E**. PBMC were stimulated and ada or control IgG was added on day 3. The level of SIRPG was analyzed on day 6. Representative overlay histograms are shown in the left panel and the cumulative results from three donors are shown in the right panel. **F**. PBMC were treated with ada or control IgG without anti-CD3 stimulation. The surface level of SIRPG in T and non-T cells was analyzed on day 3. Representative overlay histograms are shown in the left panel and cumulative results from three donors are shown in the right panel.

### Identification of SIRPG regulated genes

The function of SIRPG is still poorly understood. To investigate the function of SIRPG, we transfected anti-CD3-stimulated PBMC with control or two SIRPG sgRNAs. When examined 5 days after the transfection, more than 95% of the surviving cells were T cells, including CD4+ and CD8+ T cells (Figure 3A), and either SIRPG sgRNA reduced the expression of SIRPG in almost 80% of the CD4+ or CD8+ T cells to a level even lower than that observed in CD19+ cells of unmanipulated PBMC (Figure 3B), which expressed little or no SIRPG. The transfected T cells were maintained with IL-2 followed by re-stimulation with anti-CD3/IL-2. We found no difference in the expression of several activation markers, such as PD-1 and CD25 (Figure S3A), and the production of IFNγ and TNFα (Figure S3B) between control sgRNA-transfected (WT) and SIRPG sgRNA-transfected (KO) CD4+ or CD8+ T cells.

**Figure 3.**
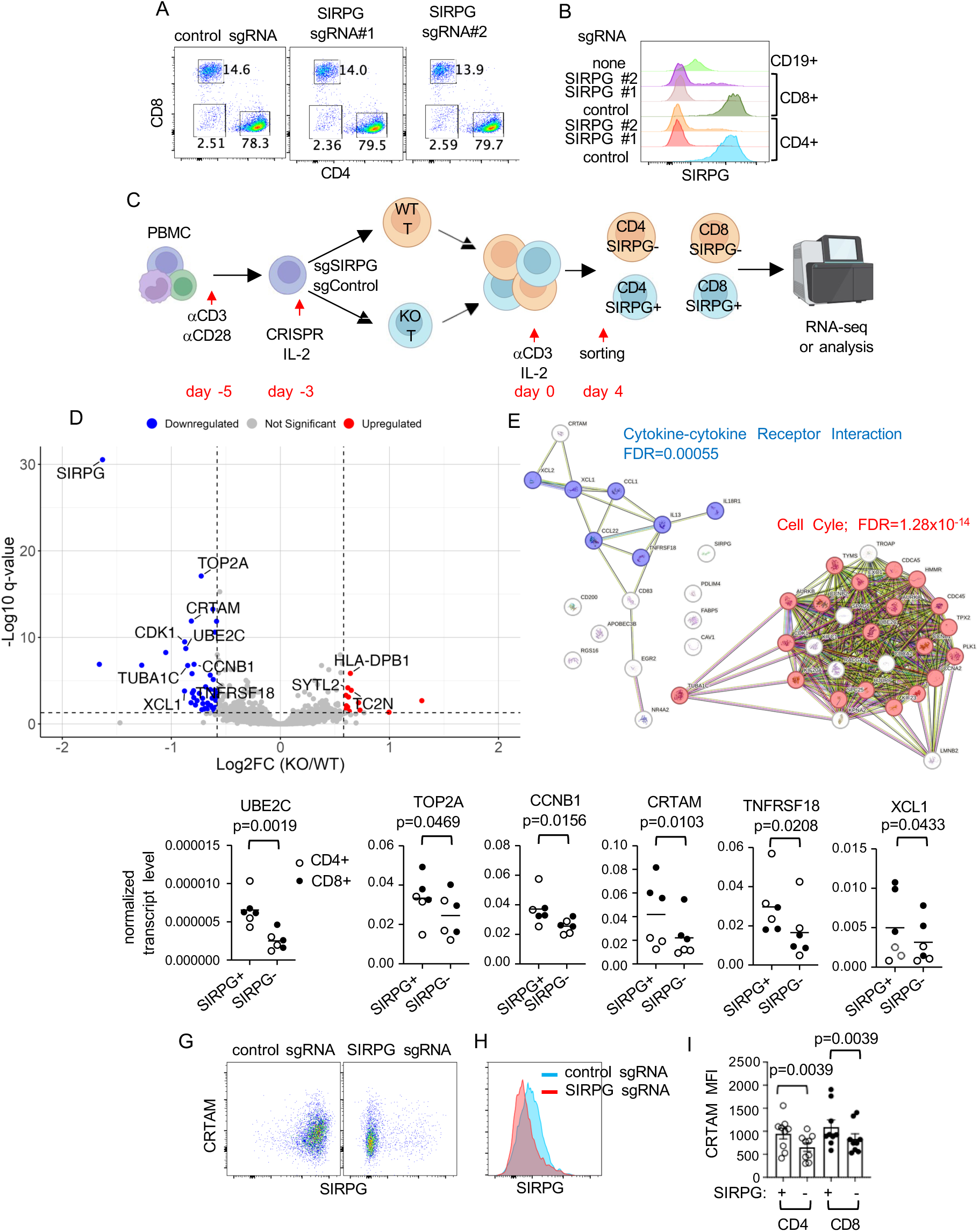
Impacts of SIRPG on the transcriptome of T cells. **A** & **B**. PBMC were stimulated and transfected with control sgRNA or two SIRPG sgRNA. The cells were analyzed with FACS on day 4. Representative CD4/CD8 plots are shown in **A**. The expression of SIRPG of transfected CD4+, CD8+, and untransfected CD19+ B cells is shown in overlay histograms (**B**). **C**-**E**. The sgRNA transfected PBMC were restimulated with anti-CD3/IL-2 for 4 days. CD4+ and CD8+ T cells were sorted into SIRPG^+^ and SIRPG^−^ populations and subjected to bulk RNA-seq (**C**). The data from sorted CD4+ and CD8+ T cells was analyzed together in a volcano plots (**D**). The genes that are down-regulated in SIRPG^−^ population were analyzed with STRING (**E**). **F**-**I**. The differential expression of indicated genes was confirmed with qPCR using independently generated/sorted CD4+ and CD8+ cells from a different set of donors (**F**). The expression of CRTAM was also examined with FACS. Representative SIRPG/CRTAM dot plots are shown in **G**; representative CRTAM overlay histograms are shown in **H**; and the CRTAM MFI from all donors is shown in **I**.

We subsequently mixed the WT and SIRPG KO cells from each donor (a total of 6 donors) at 1:1 ratio and stimulated the coculture with anti-CD3/IL-2 for 4 days, then sorted the stimulated cells into 4 populations: SIRPG^+^ CD4+, SIRPG^−^ CD4+, SIRPG^+^ CD8+ and SIRPG^−^ CD8+ (Figure 3C). The coculture enabled SIRPG^+^ and SIRPG^−^ cells to be equally stimulated. The sorted cells were then subjected to bulk RNA-seq. One SIRPG^+^ CD8 sample and its corresponding SIRPG^−^ CD8 sample were of poor quality and thus excluded from the analysis. After excluding low-expressing genes, we found a few differentially expressed genes (DEGs), including SIRPG, between SIRPG^+^ and SIRPG^−^ CD4 cells (Figure S3C, Supplemental Table 1) and between SIRPG^+^ and SIRPG^−^ CD8 cells (Figure S3D, Supplemental Table 2) based on the threshold of ≥1.5-fold change and FDR ≤0.05 in pair-wise comparison. To increase the sensitivity of detecting DEGs, we pooled SIRPG^+^ CD4 together with SIRPG^+^ CD8 and SIRPG^−^ CD4 together with SIRPG^−^ CD8. By comparing the pooled SIRPG^+^ and pooled SIRPG^−^ samples, we identified 46 genes, including SIRPG, that passed the threshold of downregulated genes in SIRPG^−^ cells compared to SIRPG^+^ cells (Figure 3D and Supplemental Table 3). STRING analysis of the down-regulated genes revealed that 20 the 46 genes, such as UBE2c, TOP2A, and CCNB1, fall into the “Cell Cycle” Reactome Pathway (FDR=1.28×10^−14^), whereas 7 of the 46 genes, including XCL1, XCL2, and CCL1, are in the “Cytokine-cytokine Receptor Interaction” KEGG Pathway (FDR=0.00055) (Figure 3E). Interestingly, several surface markers, such as CRTAM, and TNFRSF18, were also downregulated. By contrast, there are only 15 genes, including TC2N and three MHC II molecules HLA-DPB1, HLA-DRB1, and HLA-DRB5, that are upregulated in pooled SIRPG^−^ cells compared to pooled SIRPG^+^ cells (Figure 3D). We then used qPCR to confirm the differential expression of several cell cycle genes, such as UBE2C, TOP2A, and CCNB1, and surface molecules, such as CRTAM and TNFRSF18, and XCL1 between SIRPG^+^ and SIRPG^−^ cells, which were similarly generated, stimulated, and sorted in independent experiments using a different set of donors (Figure 3F). The downregulation of CRTAM in SIRPG^−^ cells was also detected with FACS (Figure 3G-3I).

### Preferential proliferation of SIRPG^+^ T cells upon activation

The strong cell cycle signal suggests that SIRPG regulates the proliferation of T cells. The transcript count of *MKI67*, which encodes the proliferation marker Ki-67, was also statistically lower in SIRPG KO cells but the fold reduction did not reach the threshold of 1.5-fold (Figure 4A). We therefore repeated CRISPR, set up the coculture, restimulated the coculture with anti-CD3/IL-2, and stained the cells for intracellular Ki-67. Indeed, the MFI of Ki-67 was already lower in SIRPG KO cells prior to restimulation compared to WT cells (Figure 4B). This difference was observed in both CD4+ and CD8+ cells and not due to less activation of SIRPG^−^ cells because the MFI of CD25 and PD-1, two activation markers, was not lower in SIRPG^−^ population compared to SIRPG^+^ population (Figure S3A). In agreement with the difference in the level of intracellular Ki-67, the number of WT cells also increased more robustly than SIRPG KO cells when separately stimulated for 4 days (Figure 4C) and the ratio of SIRPG^+^/SIRPG^−^ in the restimulated coculture increased from day 0 to day 4 (Figure 4D). This increase was comparably observed in CD4+ cells and CD8+ cells, resulting in no alteration in the CD4/CD8 ratio. Restimulation of the coculture with anti-CD3+/− IL-2 on day 4 further increased the SIRPG^+^/SIRPG^−^ ratio (Figure 4E). Thus, providing SIRPG in trans did not rescue the proliferation of SIRPG^−^ cells. The preferential expansion of SIRPG^+^ cells was not due to attenuated apoptosis because the percentages of AnnexinV^+^/7AAD^+^ cells were comparable between SIRPG^+^ and SIRPG^−^ cells (Figure S4). In addition, the preferential expansion of SIRPG^+^ cells required anti-CD3 stimulation because no such preferential expansion was observed when the coculture was maintained with only IL-2 for up to 2 weeks (Figure 4F).

**Figure 4.**
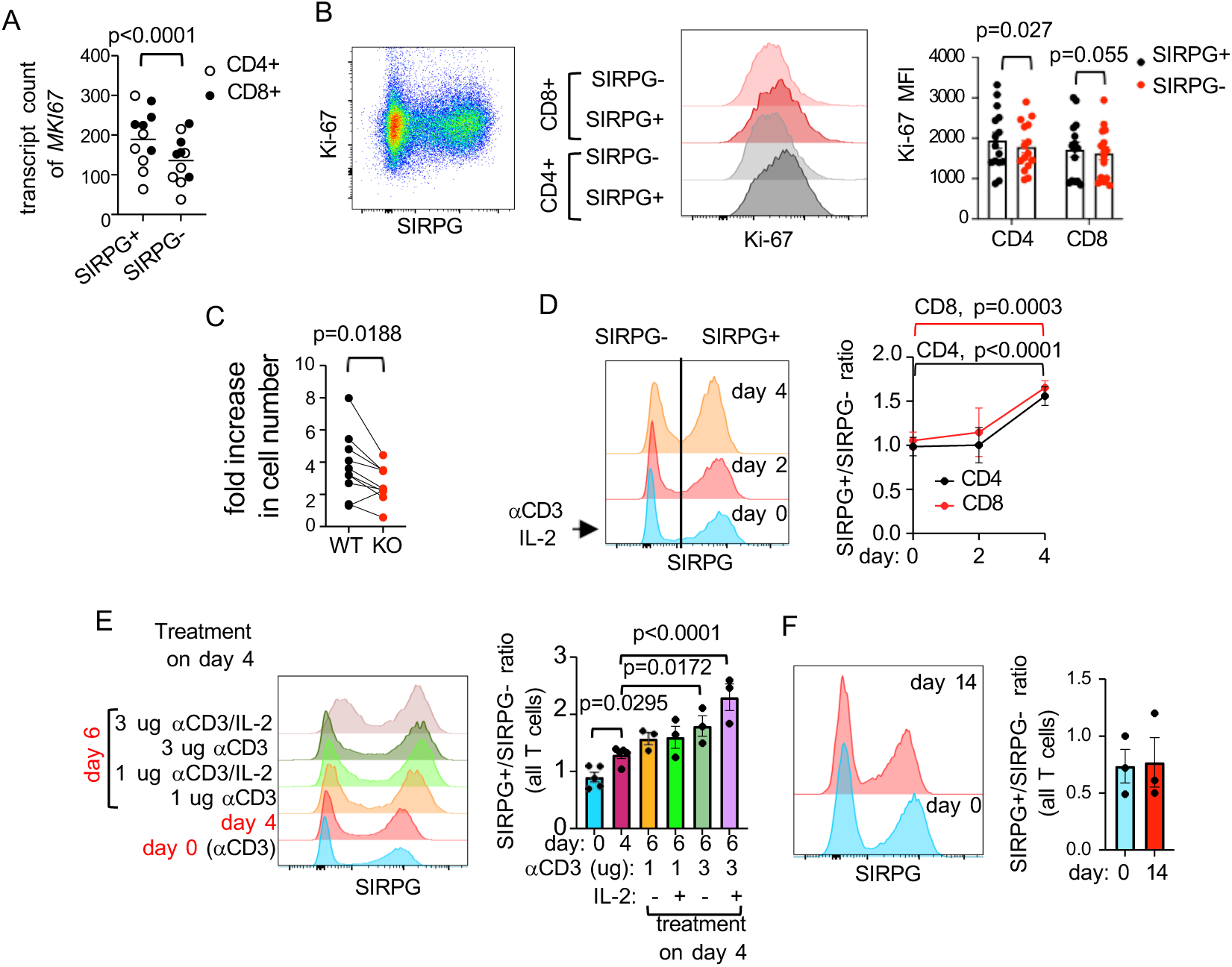
Regulation of activation-induced proliferation of T cells by SIRPG. **A**. The *MKI67* transcript counts of CD4 and CD8 T cells from the RNA-seq data shown in Figure 3 are shown. **B**-**F**. PBMC were stimulated and transfected with control sgRNA or SIRPG sgRNA. The transfected cells were mixed at 1:1 ratio. Their surface level of SIRPG and intracellular level of Ki-67 were analyzed with FACS (**B**). Representative SIRPG/Ki-67 dot plots, Ki-67 overlay histograms, and bar graphs of Ki-67 MFI from more than 6 independent donors are shown in the left panel, the middle panel, and the right panel of **B**, respectively. The transfected cells were also restimulated separately and enumerated on day 4. The fold increase in the cell number is shown in **C**. The SIRPG level of the coculture over the course of 4 days is shown in representative overlay histograms in **D** and the SIRPG^+^/SIRPG^−^ ratio from mor than 3 independent donors are shown in the right panel of **D**. A fraction of the coculture was restimulated with anti-CD3 +/− IL-2 on day 4 and analyzed on day 6. Representative overlay histograms of SIRPG are shown in the left panel of **E** and the cumulated SIRPG^+^/SIRPG^−^ ratio is shown in the right panel of **E**. Another fraction of the coculture was maintained with IL-2 only without anti-CD3 stimulation and the expression of SIRPG was analyzed on day 0 and day 14. Representative overlay histograms of SIRPG are shown in the left panel of **F** and the SIRPG^+^/SIRPG^−^ ratios on day 0 and day 14 are shown in the right panel of **F**.

### Limited role of CD47 in mediating the T cell-autonomous function of SIRPG

The only known ligand of SIRPG is CD47, which is expressed ubiquitously, including in T cells (Figure 5A). If the SIRPG-mediated proliferation and expression of CRTAM requires its interaction with CD47, then deficiency of CD47 should also lead to impaired proliferation and CRTAM expression. To examine the dependence on CD47 of SIRPG-mediated proliferation and CRTAM expression, we also generated CD47^−^ and control CD47^+^ primary T cells with CRISPR followed by sorting (Figure 5A and 5B). Ablation of CD47 had little impact on the expression of SIRPG (Figure 5B). Unlike SIRPG^−^ cells, CD47^−^ cells expanded very comparably to CD47+ cells when stimulated separately (Figure 5C). In addition, when the CD47^−^ and CD47^+^ cells were mixed at 1:1 ratio and stimulated together, the CD47^+^/CD47^−^ ratio decreased over a course of 4 days (Figure 5D), a trend consistent with the observation that murine CD47^−^ cells proliferate more robustly than CD47^+^ cells in response to anti-CD3/anti-CD28 stimulation in vitro (25). The CRTAM MFI of CD47^−^ cells was also comparable to that of CD47^+^ counterparts in the coculture (Figure 5E). These features are either different or opposite to those of SIRPG^−^ cells, suggesting that the SIRPG-mediated proliferation and expression of CRTAM is independent of CD47.

**Figure 5.**
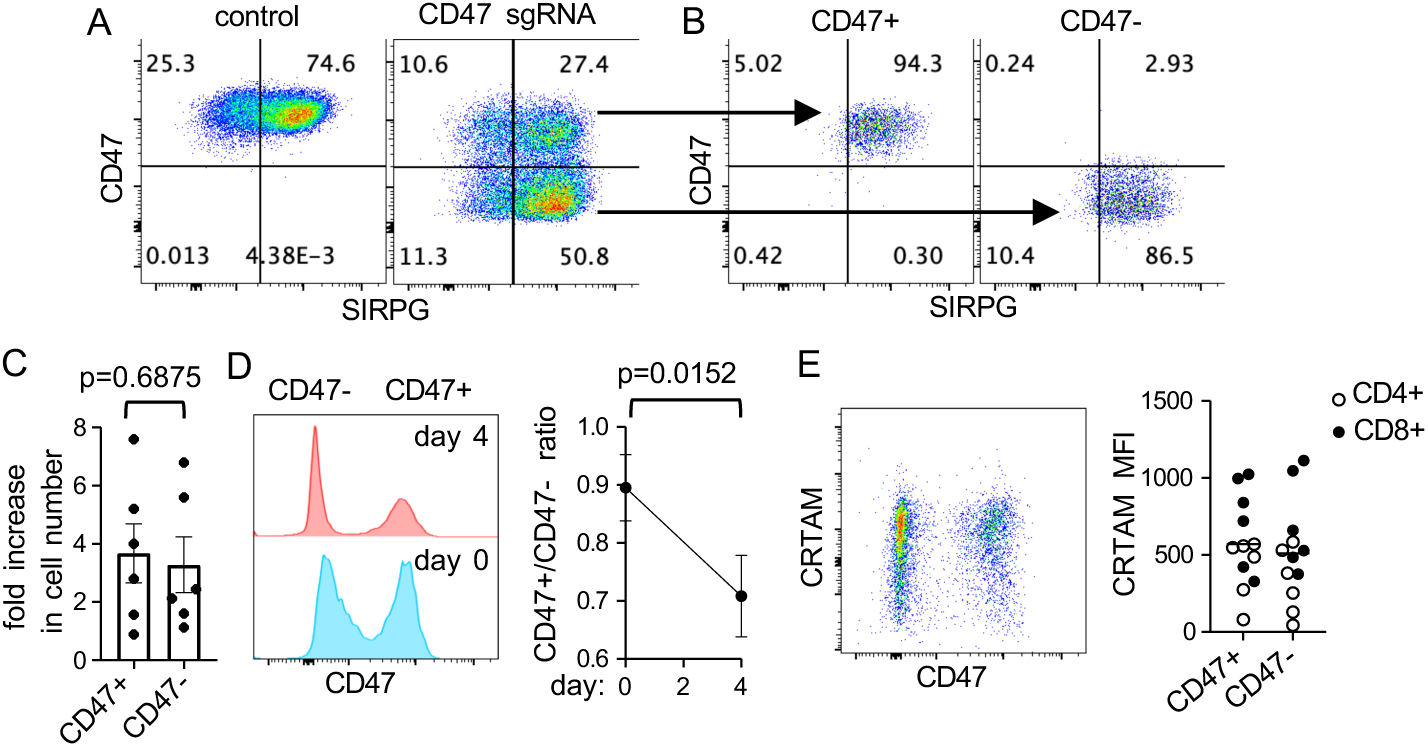
CD47 independent function of SIRPG. PBMC were transfected with control or CD47 sgRNA, analyzed with FACS for the expression of CD47 and SIRPRG 3 days later (**A**), and sorted into CD47^+^ and CD47^−^ populations (**B**). The sorted cells were re-stimulated with anti-CD3 or cocultured at 1:1 ratio before restimulation for 4 days. The fold increases in the number of separately stimulated cells are shown in **C**. Representative CD47 overlay histograms of the co-cultured cells are shown in the left panel of **D**. The CD47^+^/CD47^−^ ratio of the coculture from more than 3 donors is shown in the right panel of **D**. The expression of CRTAM of the coculture was also examined with FACS. A representative CD47/CRTAM dot plot and CRTAM MFI from all experiments (6 donors) are shown in **E**.

The SIRPG-mediated proliferation was also recapitulated in Jurkat cells, which normally express a high level of SIRPG (Figure 6A). We transfected Jurkat cells with SIRPG sgRNA. The efficiency of SIRPG sgRNA was approximately 75% when examined 3 days after the transfection (Figure 6A), resulting in de facto coculture of SIRPG^+^ and SIRPG^−^ cells. This coculture allowed us to compare their expansion without any stimulation. In agreement with the data obtained from primary T cells, the SIRPG^+^/SIRPG^−^ ratio in the SIRPG sgRNA-transfected samples increased over a course of 2-3 weeks (Figure 6A and 6B). We further sorted SIRPG^+^ and SIRPG^−^ Jurkat cells from the SIRPG sgRNA-transfected samples to more than 90% purity (Figure 6C) and co-cultured the cells at 1:1 ratio (Figure 6D). Again, the SIRPG^+^/SIRPG^−^ ratio increased from approximately 1 to nearly 3 over a course of 2 weeks (Figure 6D) and the percentage of Ki67^+^ cells and the MFI of Ki67 were lower in SIRPG^−^ cells than those in SIRPG+ cells (Figure 6E). Jurkat cells also express a high level of CD47 (Figure 6F). The efficiency of CD47 sgRNA was approximately 50%, also resulting in a de facto coculture of CD47^+^ and CD47^−^ cells. However, the ratio of CD47^+^/CD47^−^ did not increase over a course of more than 2 weeks (Figure 6F).

**Figure 6.**
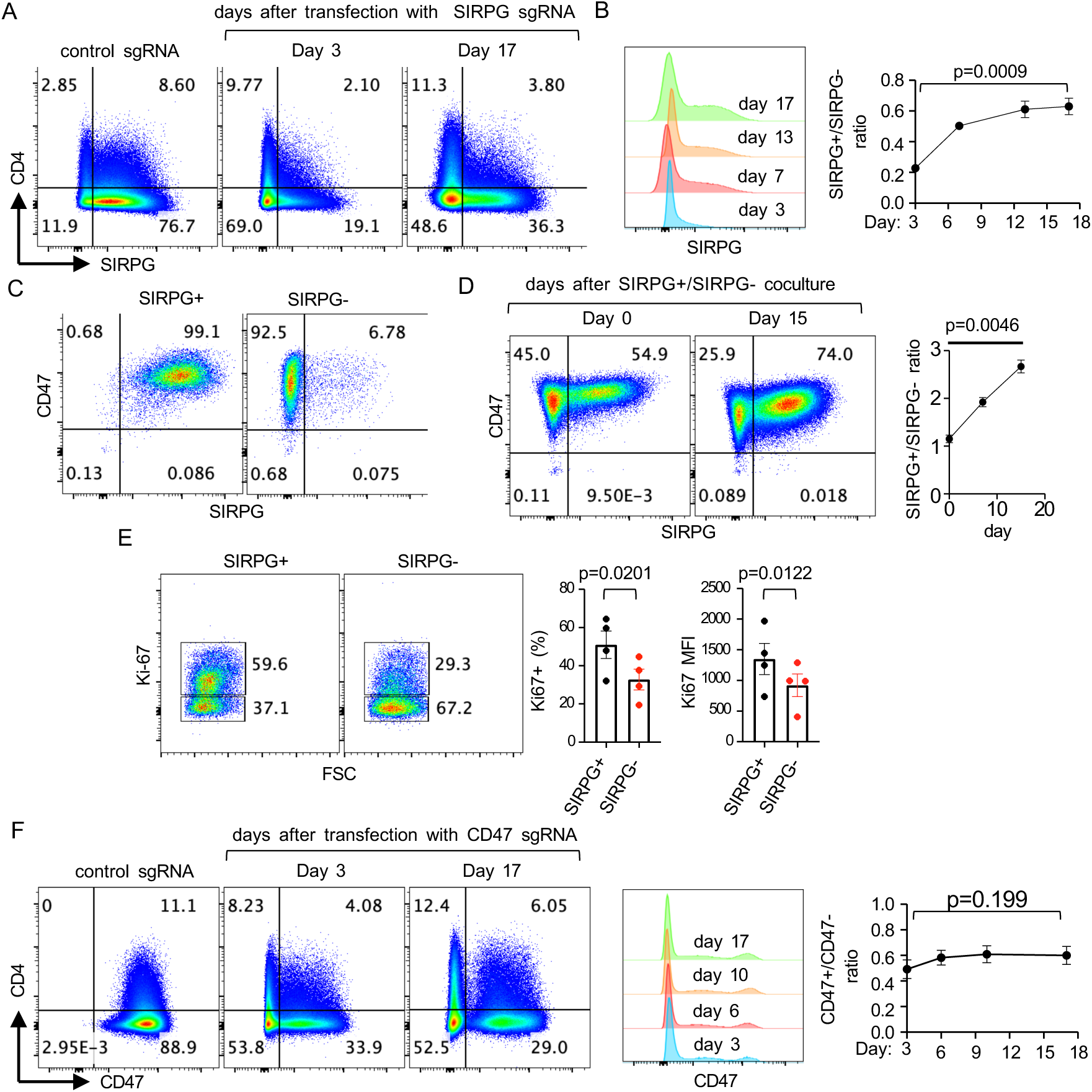
Recapitulation of the function of SIRPG in Jurkat cells. **A**-**F**. Jurkat cells were transfected with control (**A-F**), SIRPG sgRNA (**A-E**), or CD47 sgRNA (**F**) and analyzed with FACS at the indicated time points. Representative CD4/SIRPG and CD4/CD47 dot plots are shown in **A** and **F**, respectively. The expression of SIRPG and CD47 at various time points is shown in the representative overlay histograms of **B** and **F**, respectively. The SIRPG^+^/SIRPG^−^ ratio and CD47+/CD47-ratio over the time course is shown in the right panel of **B** and **F**, respectively. The SIRPG sgRNA-transfected Jurkat cells were sorted into SIRPG^+^ and SIRPG^−^ populations (**C**), cocultured at 1:1 ratio (**D** & **E**), analyzed for the expression of SIRPG (**D**), CD47 (**D**), and intracellular Ki-67 (**E**). The SIRPG^+^/SIRPG^−^ ratio of the coculture over a course of 2 weeks is shown in the right panel of **D** (at least 3 experiments). The percentage of Ki-67^+^ cells and the Ki-67 MFI are shown separately in **E**.

### Structural and functional analyses of SIRPG

To confirm the role of SIRPG in promoting proliferation, we stably expressed in KO Jurkat cells an exogenous Flag-tagged SIRPG (Flag-WT) driven by a CMV promoter (Figure 7A). While we were able to obtain nearly 100% SIRPG^+^ population after selection and sorting (Figure S5A), the percentage of SIPRG^+^ fluctuated over a wide range (20% to 98%) (Figure S5B). The cause of this fluctuation is still unclear and is probably due to the instability of the CMV promoter after integration into genome (26, 27). However, we found that the ectopic Flag-WT increased the expression of Ki-67 compared to its empty vector (Figure 7B and 7C) and that the number of Flag-WT Jurkat cells expanded more robustly than control cells over a course of one week (Figure 7D). We also used the dilution of CFSE to quantify the proliferation of Jurkat cells and found that the CFSE dilution is modestly higher in Flag-WT Jurkat cells compared to control SIRPG KO Jurkat cells (Figure S5C).

**Figure 7.**
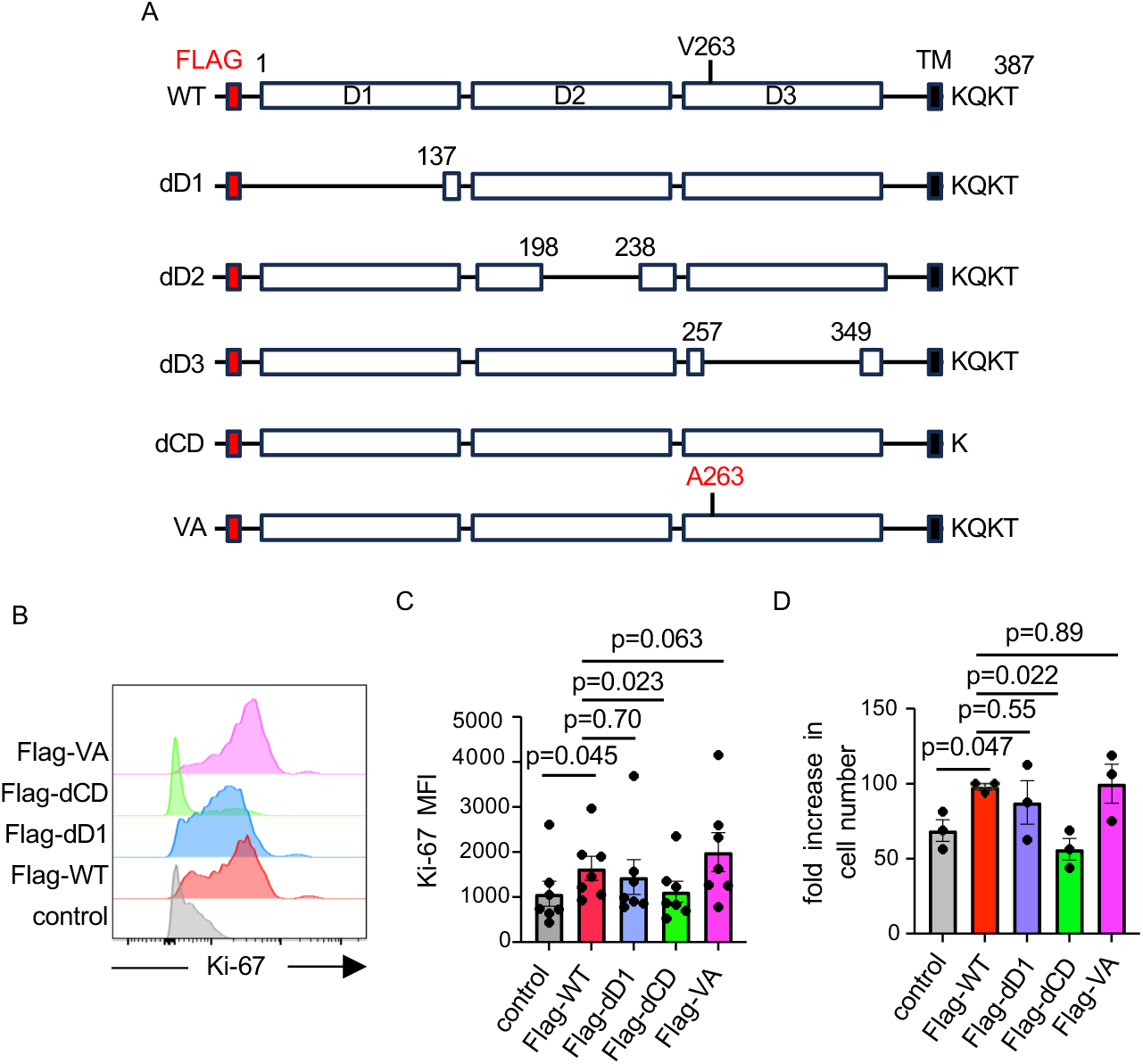
Structural and functional analysis of SIRPG. Schematic diagrams of WT Flag-SIRPG and its various deletion mutants are shown in **A**. **B**-**D**. SIRPG KO Jurkat cells stably expressing indicated Flag-SIRPG constructs were analyzed for the expression of Ki-67. Representative overlay histograms (**B**) and Ki-67 MFI (**C**) from more than 6 experiments are shown. The fold increases in the number of Jurkat cells expressing Flag-SIRPG over 7 days are shown in **D** (N=3).

We subsequently generated a series of deletion mutants of Flag-SIRPG (Figure 7A). The mutants were separately introduced into SIRPG KO Jurkat cells. While deletion of the D1 domain (dD1) or the cytoplasmic domain (dCD) did not affect the surface expression of SIRPG, deletion of the D2 (dD2) or D3 (dD3) domain rendered the Flag-SIRPG barely detectable on the surface (Figure S5A), suggesting that D2 and D3 are critical for maintaining the structure and/or stability of SIRPG protein. Again, the percentage of SIRPG^+^ cells driven by each mutant fluctuated even after selection and repeated sorting. Flag-dD1 also enhanced the expression of Ki-67, the proliferation and the expansion of Jurkat cells (Figure 7B-7D, and S5C). By contrast, Flag-dCD was unable to restore the expression of Ki-67 or cell proliferation/expansion. The V263A SNP was predicted to cause a conformational change altering the stability, structure, or function of SIRPG protein and was very likely a causal SNP of type 1 diabetes. We also introduced the SNP into the Flag-SIRPG (Flag-VA). It was as readily detected with FACS as Flag-WT 3 days after the transfection (Figure S5A), indicating that the Flag-VA mutant is structurally stable enough to be expressed on the plasma membrane. It was also as efficient as Flag-WT in restoring the expression of Ki-67 and proliferation/expansion of SIRPG KO Jurkat cells (Figure 7B-7D, and S5C). Taken together, our structural and functional analyses indicate that SIRPG-mediated proliferation depends on its CD but not its D1 and that the V263A conversion minimally influences this function.

## Discussion

The presented data reveal a novel role for SIRPG in cell-autonomously promoting activation-induced T cell proliferation and CRTAM expression. Notably, this SIRPG-driven proliferation occurs independently of CD47 but necessitates the presence of the SIRPG cytoplasmic domain (CD), implying the existence of novel SIRPG ligand and suggesting that its CD possesses signaling capabilities. These findings establish a basis for future research focused on identifying this putative ligand and elucidating the downstream signaling pathways of SIRPG. The use of CRISPR-mediated SIRPG ablation in primary T cells enhanced this study’s rigor by avoiding limitations associated with antibody or fusion protein approaches and the direct comparison of naturally occurring SIRPG^high^ and SIRPG^low^ cells. However, the absence of in vivo validation represents a current limitation.

Prior to this study, the expression profile of SIRPG on T cells in different functional states within inflamed environments was unknown. Our novel data uncovers a distinct pattern: the majority of GzmB^+^/GzmK^−^ CD8 T cells in both PBMC and inflamed synovial fluid are SIRPG^−^, whereas GzmK^+^ CD8 T cells (regardless of GzmB expression) are highly enriched within the SIRPG^+^ population in both compartments. These observations are consistent with the sc-RNAseq data from rheumatoid arthritis AMP phase 2 study (28), which showed lower SIRPG transcript levels in the CD8 GzmB^+^/Temra (T-15) cluster compared to CD8 GzmK^+^ memory (T-13 and T-14) clusters. Given that GzmB is a key cytotoxic molecule and GzmK activates the complement cascade by cleaving C2 and C4 (29, 30), and considering the enrichment of GzmK^+^ CD8 T cells in various inflamed tissues (23, 30–32) where they likely contribute to pathogenesis, the lack of a reliable surface marker for these cells has been a limitation. Our data suggests that SIRPG can serve as a valuable surface marker for identifying and enriching GzmK^+^ CD8 T cells, thus enabling future in vitro functional studies.

Despite the strong association between SIRPG and T1D, its role as pathogenic or protective remains unclear. The finding that SIRPG promotes T cell proliferation aligns with prior studies showing that SIRPG engagement with LSB2.20 enhances proliferation of anti-CD3 stimulated T cells (5), and is consistent with the T lymphopenia phenotype predicted for SIRPG by ARCHS^4^ (Massive Mining of Publicly Available RNA-seq Data from Human and Mouse) (33). This suggests a potential pathogenic role for SIRPG in T1D. However, SIRPG’s high expression on Treg cells (20, 34) introduces a layer of complexity. While it doesn’t directly impact Treg suppressive function (20), it might enhance Treg proliferation, potentially conferring protection in T1D. Interestingly, we observed that ada and eta, but not cert, augment SIRPG expression in bulk peripheral T cells, with this effect lasting for several days. Given the clinical benefits observed with TNFα inhibitors like eta and golimumab in early T1D clinical trials (35–38), future studies comparing the efficacy of different TNFα inhibitors in T1D will be crucial.

SIRPG is also expressed in tumor infiltrating lymphocytes, particularly those bearing signature of CD4+ Treg, exhausted CD8+ T, and CD8+CD103^+^ T (T_RM_) cells (34, 39, 40), and its transcript levels correlate with that of PD-1 and CTLA4 (34, 40, 41). A prior study indicated that ectopic expression of SIRPG in Jurkat cells increased PD-1 and decreased IFNg production, with opposite effects upon SIRPG knockdown (34). However, our Jurkat cells exhibit minimal PD-1 expression, preventing us from analyzing SIRPG’s impact on it. Furthermore, PD-1 and IFNγ were not significantly altered in our RNA-seq analysis or detected as differentially expressed in our SIRPG KO primary T cells. The reason for this discrepancy remains unclear, though we consider data from primary T cells more reliable than that from Jurkat cells. The normal IFNγ production in our SIRPG KO primary T cells aligns with published findings showing minimal impact of LSB2.20 on IFNγ production in anti-CD3 stimulated T cells (20). Interestingly, LSB2.20 did inhibit IFNγ production in a Vβ3^+^ T cell clone stimulated with SEE-pulsed B lymphoblastoid cells (5), suggesting that co-stimulatory signals might be necessary for SIRPG to influence IFNγ production.

Although CD47 is currently the only known ligand for SIRPG, substantial evidence points towards the existence of additional, yet unidentified, binding partners: First, the antibody KWAR23, intended to block the SIRPG-CD47 interaction, fails to inhibit SIRPG recruitment to the immune synapse, suggesting that the recruitment is independent of CD47 (20). Second, a SIRPG-CD4 fusion protein binds poorly to resting PBMCs, which naturally express high levels of CD47, but shows significantly enhanced binding to Con A-activated PBMCs, even though CD47 levels are only marginally increased (21). Moreover, our observation that the phenotypic consequences of CD47 deficiency are inconsistent with or even opposite to those of SIRPG deficiency strongly supports CD47-independent functions of SIRPG. This notion is further reinforced by the finding that the D1 domain of SIRPG, essential for CD47 binding (42), is largely unnecessary for SIRPG-mediated proliferation. The question of whether this CD47-independent function relies on another ligand remains open. However, the ability to recapitulate SIRPG’s function in isolated Jurkat cells suggests that any such ligand is likely expressed on T cells themselves.

Fuchs et al. recently defined a 16-gene signature to identify viral antigen-responsive CD8+ T cells in vitro (43). Notably, our RNA-seq analysis revealed downregulation of 7 of these 16 genes (CRTAM, TNFRSF9, EGR2, XCL2, FABP5, XCL1, and NR4A) in SIRPG KO cells, suggesting SIRPG’s involvement in regulating this viral response gene set. However, SIRPG itself was not differentially expressed in Fuchs’ study, the reason for which remains unclear. One compelling explanation is that viral antigen-responsive T cells might upregulate a yet-to-be-identified SIRPG ligand, rather than SIRPG, in T cells or non-T cells, triggering the expression of this gene set through T cell-T cell or T cell-other cell interactions. Furthermore, it is highly likely that this novel SIRPG ligand is expressed at considerably higher levels on non-T cells, such as antigen-presenting cells or tissue mesenchymal cells. This suggests that the CD47-independent functions of SIRPG may be particularly relevant when T cells engage with these ligand-expressing cells.

Since the D1 domain is almost unnecessary for SIRPG-driven proliferation, the novel ligand of SIRPG is highly likely to engage the D2 and/or D3 domains. The T1D-associated V263A single nucleotide polymorphism (SNP) resides within the D3 domain, suggesting a potential impact on this novel ligand interaction. However, the natural occurrence of alanine at position 263 in most non-human primate SIRPG, and its minimal effect on SIRPG-mediated proliferation in our assays, argues against this scenario. Despite this, we cannot rule out a critical role for the D3 domain in the proliferation process or the possibility that the V263A SNP affects other, yet undiscovered functions of SIRPG. The SIRPG CD, with its short KQKT sequence and the absence of a DAP12-interacting lysine in its transmembrane region (unlike SIRPB), has led to the widely held belief that SIRPG lacks intrinsic signaling capacity. Our data, showing an absolute requirement for the SIRPG CD in mediating proliferation, provides the first direct evidence supporting a critical role for this domain in signal transduction. The precise mechanism by which the SIRPG CD conveys proliferative signals, however, remains to be elucidated. Future studies employing site-specific mutagenesis targeting D2, D3, and each residue within the CD will be essential to define the structural determinants of SIRPG in supporting T cell proliferation.

The highly restricted expression of SIRPG, predominantly in primate T cells, strongly suggests a crucial role in adaptive immunity and positions it as a promising therapeutic target for a range of human diseases. Our data have provided valuable new information regarding SIRPG expression and function; however, future investigations using humanized mice will be indispensable for fully elucidating its in vivo function and underlying mechanisms, thereby unlocking its therapeutic potential.

## Methods

### Human subjects

Health donor PBMCs were purified from leukoreduction collars obtained from the Crimson Biomaterials Collection Core Facility, which prospectively collects discarded clinical materials matching investigator-defined criteria against available information on clinical samples. Synovial fluid was obtained from one patient with seronegative rheumatoid arthritis and one patient with undifferentiated inflammatory arthritis at the Brigham and Women’s Hospital Arthritis Center. This study has been approved by Partners Human Research Committee (PHRC), Boston, MA.

### Preparation, culture and stimulation of synovial fluid cells and PBMCs

PBMCs were isolated from leukoreduction collars by Ficoll-Paque PLUS (17-1440-03, GE Healthcare, Pittsburgh, PA) density gradient centrifugation and cryopreserved prior to use. PBMCs were cultured in T cell media consisting of Roswell Park Memorial Institute (RPMI)-1640 supplemented with 10% heat-inactivated fetal bovine serum (FBS, BenchMark™, Gemini Bio, Sacramento, CA, USA), 1% penicillin-streptomycin (Gibco®, Waltham, MA, USA), 10 mM HEPES (Gibco®), 2mM L-glutamine (Gibco®) and 1mM sodium pyruvate (Gibco®). The cells were maintained in an incubator at 37°C, 5% CO_2_. PBMCs were plated in 24-well plates (2-2.5 millions/1ml/well) pre-coated with anti-CD3 (2,5ug/ml, OKT3, Supplemental Table 3) and soluble anti-CD28 (2,0ug/ml, CD28.2, Supplemental Table 3) and stimulated with adalimumab (125ug/ml), etanercept (50ug/ml), certolizumab pegol (50ug/ml), tocilizumab (100ug/ml), or an isotype control anti-human IgG1 (125ug/ml, #BE0297, BioXCell, Lebanon, NH, USA) for indicated times before analyzing by flow cytometry. Synovial fluid was processed by density centrifugation using Ficoll-Paque Plus (GE Healthcare) to isolate mononuclear cells. Cells were then cryopreserved for batched analysis. Cryopreserved mononuclear cells were thawed into warm RPMI-1640 with 10% fetal bovine serum (FBS, HyClone) and washed once in cold RPMI-1640 plus 10% FBS prior to antibody staining.

### RNA Sequencing and Differential Gene Expression Analysis

PBMCs were activated for two or four days and CD4⁺ and CD8⁺ T cells were sorted using fluorescence-activated cell sorting (FACS) on a BD FACS Aria Fusion sorter and collected directly into buffer TCL 2× (Qiagen, #1070498) supplemented with 1% β-mercaptoethanol for immediate cell lysis and RNA stabilization and lysates were stored at −80°C until further processing. RNA-seq libraries were prepared and processed using the SmartSeq2 protocol. Paired-end sequencing was performed on an Illumina sequencing platform (Illumina, San Diego, CA). Raw sequencing reads in FASTQ format were processed using fastp for quality control and adapter trimming. Transcript quantification was performed using Salmon (v1.3.2) in quasi-mapping mode against the reference transcriptome (GRCh38.p14). Differential gene expression analysis was conducted using the DESeq2 (v1.46.0) package in R. Genes with a fold change of ±1.5 and an adjusted p-value < 0.05 were considered significantly differentially expressed. Gene Ontology (GO) biological process enrichment analyses were conducted using the clusterProfiler (v4.10.0) R package. Pathways with an adjusted p-value < 0.05 were considered significantly enriched and Protein-protein interaction analysis was performed using the STRING database (v12.0).

### Data Availability

The RNA-seq data generated in this study have been submitted to the NCBI Gene Expression Omnibus (GEO) and are currently under revision. The dataset will be publicly accessible upon acceptance of the submission. The data are also available upon request.

### Quantitative RNA analysis

RNA isolation, reverse transcription, and quantitative PCR (qPCR) were performed as previously described (44). Gene expression levels were normalized to GAPDH from the same sample. Primer sequences used for qPCR are listed in Supplemental Table 2.

### FACS

For extracellular staining, cells were resuspended in MACS buffer (phosphate-buffered saline [1×], 1% bovine serum albumin [Sigma-Aldrich, #A9647], 2 mM EDTA) and incubated with fluorochrome-conjugated antibodies for 30 minutes at 4°C (see Supplemental Table 4 for antibody details). Synovial cells were first stained with Fixable Viability Dye eFluor 455UV (Thermo Fisher Scientific) in phosphate-buffered saline (PBS) for 20 minutes on ice before incubation with fluorochrome-conjugated antibodies. For intracellular staining, cells were first fixed and permeabilized using the BD Cytofix/Cytoperm kit (#554714) according to the manufacturer’s instructions, followed by antibody staining. Stained cells were acquired using a FACSCanto or LSRFortessa flow cytometer (Becton Dickinson, Franklin Lakes, NJ), and data were analyzed with FlowJo software (Becton Dickinson). For apoptosis analysis, cells were resuspended in Annexin V Binding Buffer (BioLegend, #422201) and incubated with APC-conjugated Annexin V (BioLegend, #640920) and 7-AAD (BioLegend, #420403) for 15 minutes at room temperature in the dark before analysis. For carboxyfluorescein succinimidyl ester (CFFSE) staining, cells were labeled with 50 nM CFSE (Thermo Fisher Scientific, #C34554) in pre-warmed PBS (37°C) at a density of 1×10⁶ cells/mL and incubated for 15 minutes at 37°C. The reaction was quenched with five volumes of cold complete T cell medium and incubated on ice for 5 minutes. Cells were then washed twice with T cell medium and cultured at 37°C, 5% CO₂ for up to 2 days. CFSE dilution was assessed by flow cytometry.

### CRISPR of human PBMC and Jurkat cells

RNP complexes were assembled with Cas9-NLS purified protein from the QB3 MacroLab (UC Berkeley) and either scramble RNA or SIRPG-specific sgRNAs (Synthego, Redwood City, CA) with the sequence GUGACCAACAGGAGCUUCUC (cut site: 1,649,368). Cells were washed and resuspended in Nucleofector™ Set P3 Solution for primary cells (#V4XP-3032, Lonza, Basel, Switzerland), and nucleofection was performed using pulse code EO-115 on a 4D-Nucleofector® (Lonza). Following nucleofection, cells were rested for 15 minutes in warm T cell medium before supplementation with 50 U/mL IL-2 (teceleukin, NIH Repository, Frederick, MD) for three days, followed by further restimulation. Jurkat cells were resuspended in SE Cell Line 4D-Nucleofector™ Solution (Cat. #V4XC-1032, Lonza) and nucleofected with control or SIRPG RNPs using pulse code CL-120.

### Cell lines, plasmid, and mutagenesis

Jurkat cells were maintained in RPMI medium containing 10% heat-inactivated fetal bovine serum (Hyclone, GE Healthcare Life Science, Marlborough, MA). A modified cDNA fragment encoding full-length human SIRPG with an N-terminal Flag was synthesized and inserted into the HindIII and NheI of pTwist-CMV-Hygro (Twist Bioscience, South San Francisco, CA). Additional Kpn1, BamH1, and Xho1 sites (one of each) were introduced into the cDNA, enabling separate deletion of D1 (with Kpn1 digestion), D2 (with BamH1 digestion), or D3 (with Xho1 digestion) domain. Site specific mutagenesis was performed with QuikChange XL Site-Directed Mutagenesis kit (Agilent Technologies, Lexington, MA) according to manufacturer’s instructions.

### Statistical analysis

Statistical analysis was performed with one-way ANOVA followed by multiple comparisons for Figure 2B, 2C, 4E, 7, and S5C, paired two-tailed t test for Figure 2D, 2E, 3, 4A, 4B, 4C, 4D, 5, and 6).

