## Supplemental Materials for "Enhancement of activation-induced T cell proliferation by SIRPG in a CD47-independent manner"

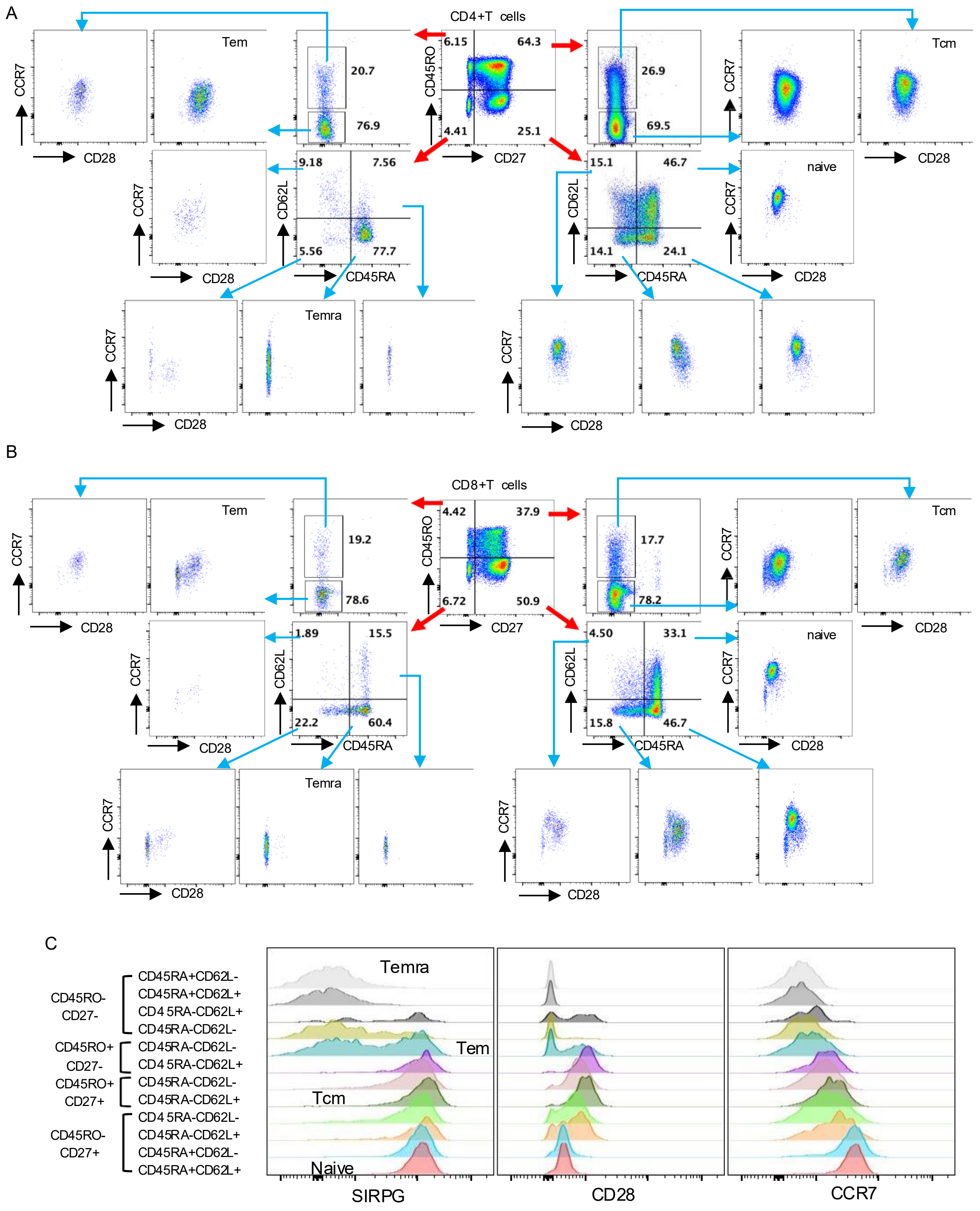


**Figure S1. The expression of SIRPG in resting peripheral blood T cells cells.** Peripheral blood CD4+ T cells (**A**), and CD8+ T cells (**B** & **C**) were stained with antibodies against indicated markers. The gating of various subsets of is shown in **A** and **B**. The overlay histograms of SIRPG, CD28, and CCR7 of various subsets of CD8+ T cells are shown in **C**.

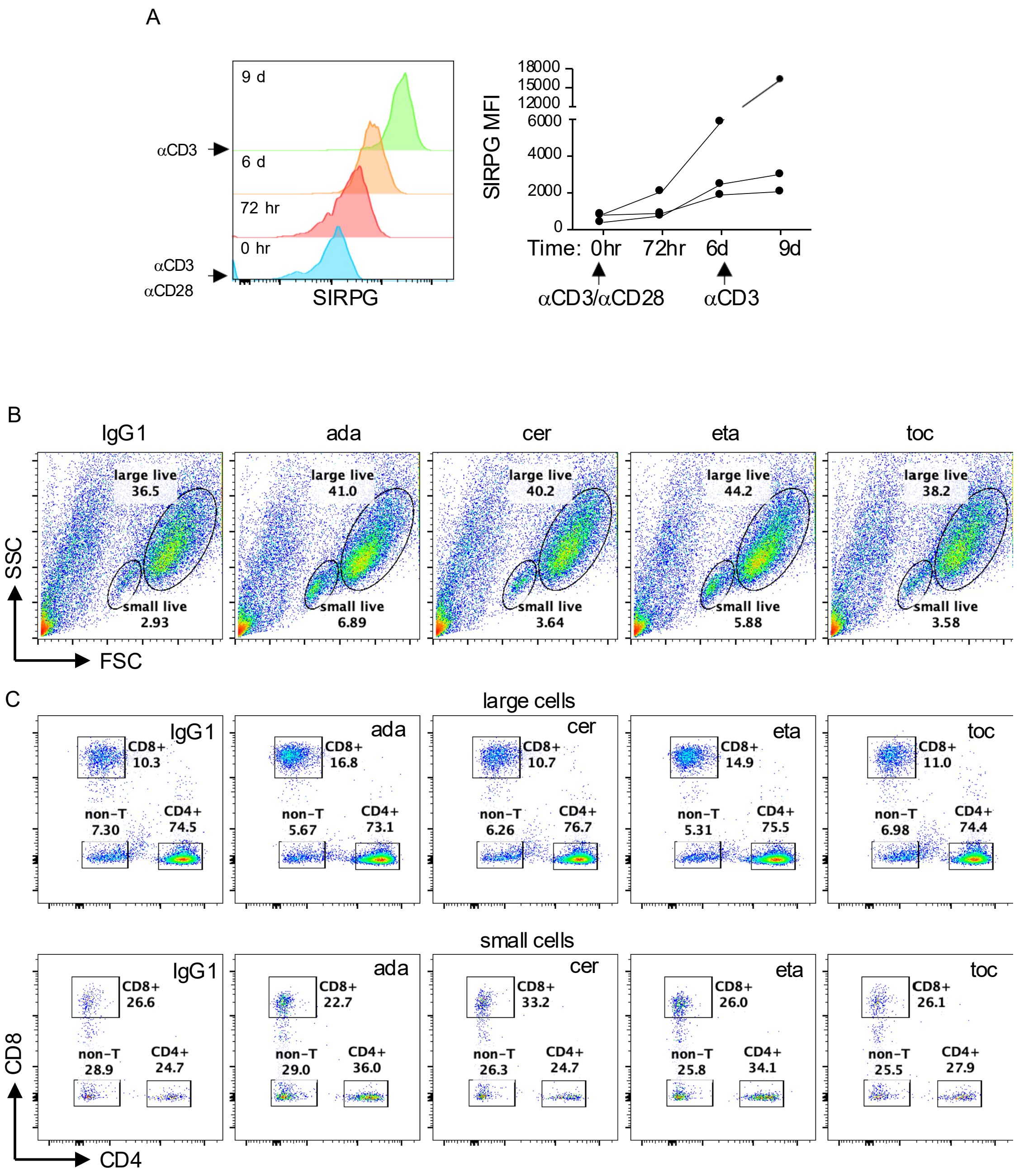


**Figure S2. Differential effects of TNFα inhibitors on the expression of SIRPG. A.** PBMC were stimulated with anti-CD3/anti-CD28 and anti-CD3 at indicated time points. The expression of SIRPG in CD8+ T cells was examined with FACS. Representative SIRPG overlay histograms are shown in the left panel and the SIRPG MFI from 3 donor is shown in the right panel. **B** & **C**. PBMC were stimulated with anti-CD3/anti-CD28 in the presence of indicated monoclonal antibodies for 3 days and analyzed with FACS. Representative FSC/SSC plots are shown in **B**; representative CD4/CD8 plots of blasting (large) and non-blasting (small) cells are shown in **C**.

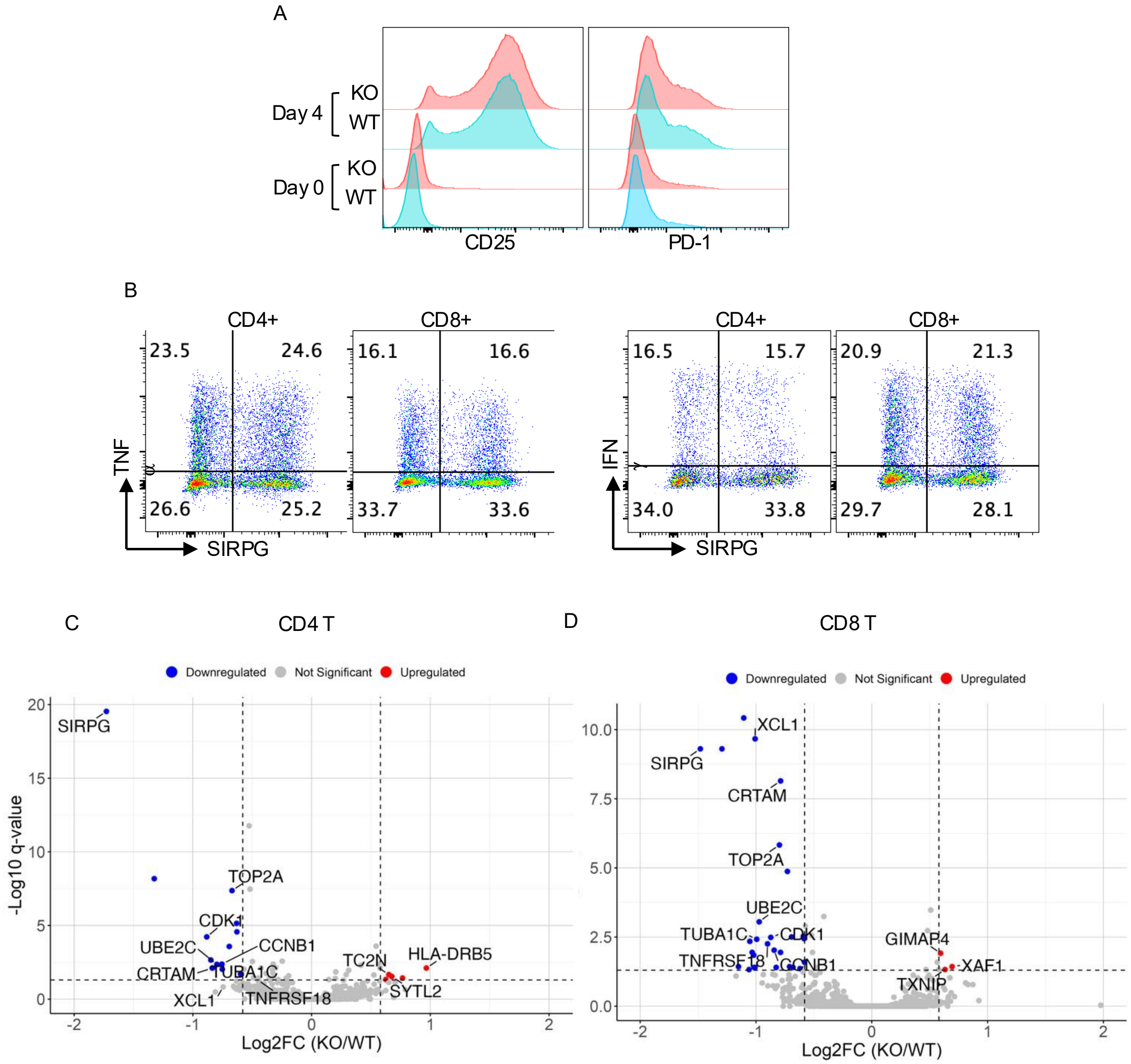


**Figure S3. Identification of SIRPG-regulated genes. A & B.** Control (WT) or SIRPG sgRNA (KO) transfected PBMC were stimulated with anti-CD3 for 4 days. The upregulation of CD25 and PD-1 was examined with FACS (**A**). The WT and SIRPG KO cells were also mixed at 1:1 ratio, stimulated with anti-CD3, and subjected to intracellular staining for TNF$\alpha$ and IFN$\gamma$ (**B**). The differentially expressed genes between SIRPG+ and SIRPG- CD4+ T cells and between SIRPG+ and SIRPG- CD8+ T cells are shown in the volcano plot of **C** and **D**, respectively.

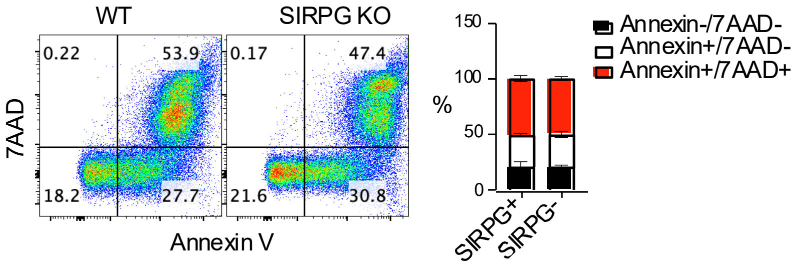


**Figure S4. Little impact of SIRPG on the apoptosis of T cells.** Stimulated WT and SIRPG KO T cells shown in Figure 4D were stained for the level of AnnexinV and 7AAD. Representative dot plots are shown in the left panel and the the percentage of indicated populations from three donors is shown in the right panel.

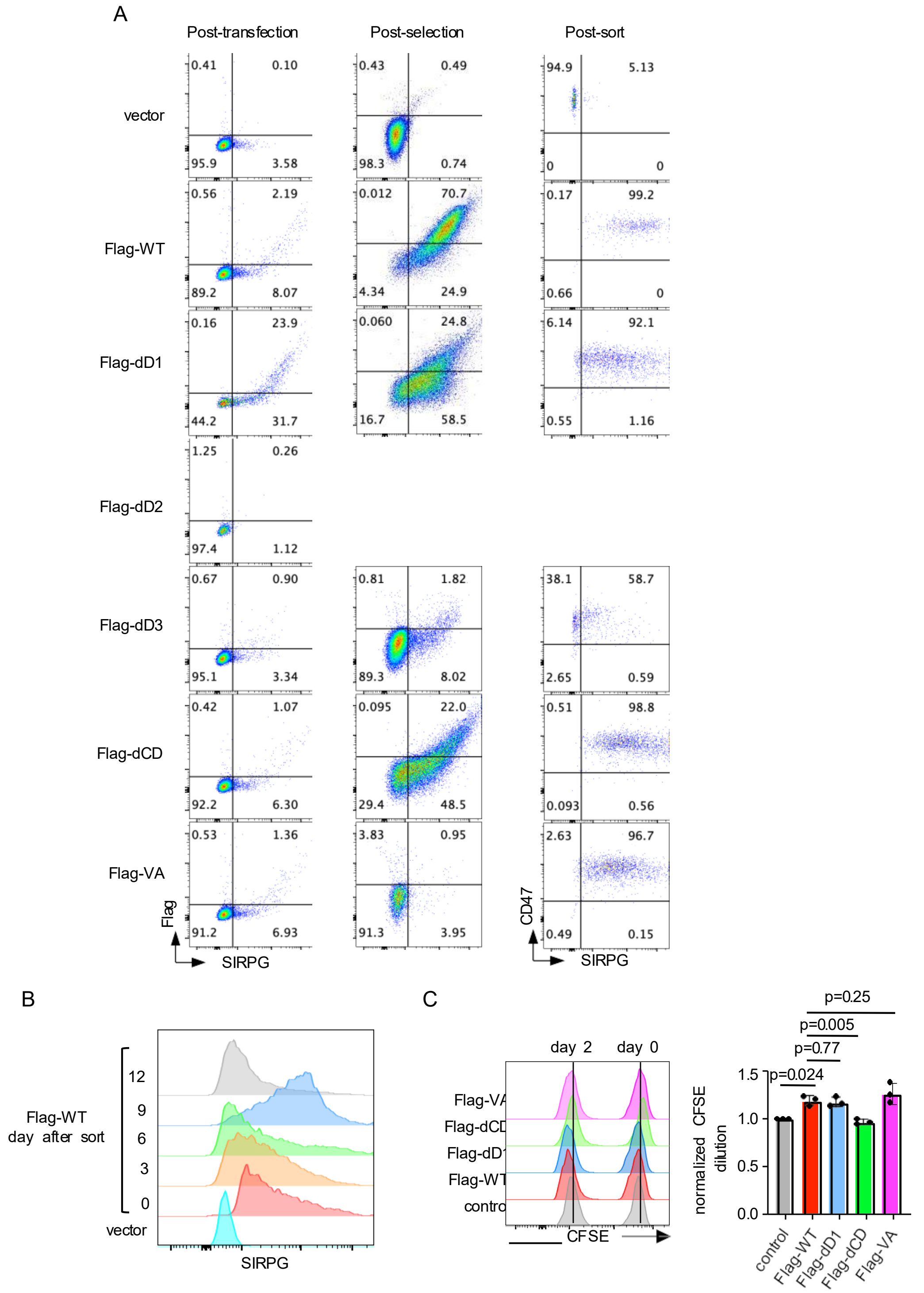


**Figure S5. Structural and functional analyses of SIRPG. A-C.** SIRPG KO Jurkat cells were transfected with indicate Flag-SIRPG constructs shown in Figure 7A. The expression of Flag-SIRPG was examined 3 days after the transfection, 3 weeks after hygromycin selection, and immediately after sorting (**A**). The expression of Flag-WT at different time points after sorting is shown in overlay histograms (**B**). The sorted cells were stained for CFSE and the level of CFSE was examined on day 0 and day 2 (the left panel of **C**). The normalized CFSE dilution over the two-day course from three independent experiments is shown in the right panel of **C**. The fold of CFSE dilution in control cells (ranging from 12 to 36) was arbitrarily set as 1.

Supplementary Table 1: Differentially expressed genes between SIRPG knockout (KO) and wild-type (WT) $CD4^+$ T cells.

| symbol | baseMean | log2FoldChange | lfcSE | stat | pvalue | padj |
|---|---|---|---|---|---|---|
| HLA-DRB5 | 59.78411 | 0.966898 | 0.238051 | 4.061726 | 4.87E-05 | 0.007745 |
| TC2N | 93.46348 | 0.650992 | 0.174647 | 3.727482 | 0.000193 | 0.021707 |
| SYTL2 | 73.50878 | 0.676196 | 0.186249 | 3.630602 | 0.000283 | 0.029163 |
| HLA-DPB1 | 203.5102 | 0.766374 | 0.216427 | 3.541022 | 0.000399 | 0.037097 |
| OASL | 91.39674 | 0.625633 | 0.182334 | 3.431247 | 0.000601 | 0.04679 |
| SIRPG | 140.3094 | -1.7283 | 0.171662 | -10.0681 | 7.65E-24 | 2.92E-20 |
| CCL22 | 98.55052 | -1.32521 | 0.192077 | -6.89938 | 5.22E-12 | 6.64E-09 |
| TOP2A | 282.0382 | -0.67017 | 0.102253 | -6.554 | 5.6E-11 | 4.28E-08 |
| EGR2 | 215.3408 | -0.63162 | 0.110617 | -5.71003 | 1.13E-08 | 7.18E-06 |
| ZBED2 | 241.6874 | -0.62887 | 0.115326 | -5.45297 | 4.95E-08 | 2.7E-05 |
| CDK1 | 104.1192 | -0.88308 | 0.167041 | -5.28662 | 1.25E-07 | 5.94E-05 |
| SEMA4A | 130.0913 | -0.69335 | 0.139731 | -4.96199 | 6.98E-07 | 0.000266 |
| CCL1 | 260.3065 | -0.84869 | 0.188032 | -4.51357 | 6.37E-06 | 0.002211 |
| CCNB1 | 96.33368 | -0.75533 | 0.175337 | -4.30789 | 1.65E-05 | 0.004193 |
| UBE2C | 77.83421 | -0.79782 | 0.186231 | -4.28402 | 1.84E-05 | 0.004378 |
| CD200 | 59.46942 | -0.83705 | 0.205598 | -4.0713 | 4.68E-05 | 0.007745 |
| CRTAM | 94.73216 | -0.83434 | 0.205193 | -4.06611 | 4.78E-05 | 0.007745 |
| TUBA1C | 84.31008 | -0.75188 | 0.187709 | -4.00554 | 6.19E-05 | 0.00939 |
| TPX2 | 106.2511 | -0.59807 | 0.158616 | -3.77053 | 0.000163 | 0.019426 |

Supplementary Table 2**:** Differentially expressed genes between SIRPG knockout (KO) and wild-type (WT) CD8$^{+}$ T cells.

| symbol | baseMean | log2FoldChange | lfcSE | stat | pvalue | padj |
|---|---|---|---|---|---|---|
| GIMAP4 | 215.5172 | 0.594756 | 0.148682 | 4.000184 | 6.33E-05 | 0.012228 |
| XAF1 | 134.1174 | 0.693915 | 0.189259 | 3.666492 | 0.000246 | 0.037116 |
| TXNIP | 130.8879 | 0.632712 | 0.178213 | 3.550316 | 0.000385 | 0.047361 |
| XCL2 | 446.339 | -1.10529 | 0.142245 | -7.77033 | 7.83E-15 | 3.78E-11 |
| XCL1 | 506.1353 | -1.00745 | 0.135113 | -7.45635 | 8.9E-14 | 2.15E-10 |
| SIRPG | 144.3338 | -1.48033 | 0.203503 | -7.27425 | 3.48E-13 | 4.96E-10 |
| CCL1 | 210.1461 | -1.29386 | 0.178413 | -7.25206 | 4.1E-13 | 4.96E-10 |
| CRTAM | 413.4742 | -0.7874 | 0.11496 | -6.84932 | 7.42E-12 | 7.17E-09 |
| TOP2A | 395.6591 | -0.7975 | 0.132677 | -6.01087 | 1.85E-09 | 1.49E-06 |
| KPNA2 | 333.735 | -0.72861 | 0.129715 | -5.61704 | 1.94E-08 | 1.34E-05 |
| UBE2C | 103.0297 | -0.97202 | 0.203792 | -4.76968 | 1.85E-06 | 0.000891 |
| EGR2 | 271.7175 | -0.5911 | 0.132796 | -4.45123 | 8.54E-06 | 0.002946 |
| TUBB4B | 229.0424 | -0.6924 | 0.156542 | -4.42313 | 9.73E-06 | 0.003132 |
| CDK1 | 154.6944 | -0.87191 | 0.198119 | -4.40096 | 1.08E-05 | 0.003253 |
| RRM2 | 468.2425 | -0.58502 | 0.134024 | -4.36502 | 1.27E-05 | 0.003611 |
| TUBA1C | 87.56303 | -0.99333 | 0.228813 | -4.34121 | 1.42E-05 | 0.003802 |
| AURKA | 73.70983 | -1.05291 | 0.24539 | -4.29076 | 1.78E-05 | 0.004526 |
| TNFRSF18 | 96.11126 | -0.90015 | 0.212658 | -4.23286 | 2.31E-05 | 0.005572 |
| CCNB1 | 128.1495 | -0.84301 | 0.205557 | -4.10108 | 4.11E-05 | 0.009458 |
| CDC20 | 155.1565 | -0.78789 | 0.195631 | -4.02743 | 5.64E-05 | 0.011348 |
| RGS16 | 71.68637 | -1.03311 | 0.255506 | -4.0434 | 5.27E-05 | 0.011348 |
| PDLIM4 | 76.23234 | -1.01915 | 0.257739 | -3.95418 | 7.68E-05 | 0.014267 |
| TNFRSF9 | 280.3794 | -0.58078 | 0.152881 | -3.79891 | 0.000145 | 0.025701 |
| TPX2 | 139.1278 | -0.71002 | 0.193029 | -3.6783 | 0.000235 | 0.037116 |
| CCL22 | 140.6534 | -2.19652 | 0.59861 | -3.66936 | 0.000243 | 0.037116 |
| CD83 | 44.60081 | -1.15075 | 0.312864 | -3.67812 | 0.000235 | 0.037116 |
| RACGAP1 | 66.79175 | -1.00827 | 0.278598 | -3.61909 | 0.000296 | 0.039665 |
| FABP5 | 184.4237 | -0.68046 | 0.187926 | -3.62088 | 0.000294 | 0.039665 |
| AURKB | 79.64339 | -0.82549 | 0.227499 | -3.62854 | 0.000285 | 0.039665 |
| APOBEC3B | 89.25181 | -1.01529 | 0.27916 | -3.63695 | 0.000276 | 0.039665 |
| TUBA1B | 1758.407 | -0.61996 | 0.173194 | -3.57957 | 0.000344 | 0.043744 |
| CD72 | 43.39631 | -1.05958 | 0.299962 | -3.53238 | 0.000412 | 0.047361 |

Supplementary Table 3: Differentially expressed genes between SIRPG knockout (KO) and wild-type (WT) including both CD4+ and CD8+ T cells.

| symbol | baseMean | log2FoldChange | lfcSE | stat | pvalue | padj |
|---|---|---|---|---|---|---|
| HLA-DPB1 | 207.0103 | 0.641926 | 0.11056 | 5.806144 | 6.39E-09 | 1.44E-06 |
| SYTL2 | 83.20699 | 0.61851 | 0.122724 | 5.039855 | 4.66E-07 | 6.64E-05 |
| TC2N | 79.07288 | 0.65012 | 0.133006 | 4.887905 | 1.02E-06 | 0.000129 |
| OASL | 91.97102 | 0.601569 | 0.132592 | 4.537002 | 5.71E-06 | 0.00049 |
| RGS1 | 104.4287 | 0.624296 | 0.141178 | 4.422058 | 9.78E-06 | 0.000772 |
| PLEK | 47.05728 | 1.296842 | 0.311173 | 4.167593 | 3.08E-05 | 0.002044 |
| HLA-DRB5 | 69.53112 | 0.71695 | 0.17816 | 4.024192 | 5.72E-05 | 0.003506 |
| IL7R | 71.10164 | 0.605254 | 0.161117 | 3.756607 | 0.000172 | 0.007889 |
| TRG-AS1 | 41.6858 | 0.616783 | 0.169274 | 3.643698 | 0.000269 | 0.011139 |
| PDE4B | 38.80476 | 0.604913 | 0.173555 | 3.485429 | 0.000491 | 0.017725 |
| TRGC2 | 38.93226 | 0.730415 | 0.218617 | 3.34107 | 0.000835 | 0.025805 |
| HLA-DRB1 | 207.4469 | 0.632239 | 0.195641 | 3.231633 | 0.001231 | 0.03377 |
| NBPF26 | 53.13375 | 0.996015 | 0.317954 | 3.132579 | 0.001733 | 0.043837 |
| SIRPG | 141.7013 | -1.63075 | 0.132064 | -12.3482 | 4.98E-35 | 2.91E-31 |
| TOP2A | 325.722 | -0.72594 | 0.076646 | -9.47139 | 2.76E-21 | 8.07E-18 |
| EGR2 | 236.8204 | -0.61897 | 0.073522 | -8.4188 | 3.8E-17 | 5.56E-14 |
| CRTAM | 218.7413 | -0.81642 | 0.101878 | -8.01372 | 1.11E-15 | 1.3E-12 |
| ZBED2 | 314.0547 | -0.58544 | 0.073323 | -7.98439 | 1.41E-15 | 1.38E-12 |
| KPNA2 | 266.783 | -0.60033 | 0.07901 | -7.5981 | 3.01E-14 | 2.51E-11 |
| CDK1 | 123.5665 | -0.8777 | 0.121229 | -7.24006 | 4.49E-13 | 3.28E-10 |
| UBE2C | 87.48298 | -0.86732 | 0.124271 | -6.97922 | 2.97E-12 | 1.93E-09 |
| CCL1 | 240.1543 | -1.05099 | 0.154247 | -6.8137 | 9.51E-12 | 5.56E-09 |
| CCL22 | 114.7806 | -1.66006 | 0.264603 | -6.27377 | 3.52E-10 | 1.29E-07 |
| CCNB1 | 108.4851 | -0.79172 | 0.126105 | -6.27827 | 3.42E-10 | 1.29E-07 |
| XCL2 | 190.6097 | -1.27163 | 0.204212 | -6.227 | 4.75E-10 | 1.63E-07 |
| TUBA1C | 85.37743 | -0.84997 | 0.137155 | -6.19718 | 5.75E-10 | 1.77E-07 |
| CD200 | 62.7888 | -0.81032 | 0.139997 | -5.7881 | 7.12E-09 | 1.54E-06 |
| TPX2 | 118.7997 | -0.64339 | 0.112584 | -5.71477 | 1.1E-08 | 2.29E-06 |
| TNFRSF18 | 156.0165 | -0.61551 | 0.111756 | -5.50761 | 3.64E-08 | 7.33E-06 |
| AURKB | 67.30321 | -0.68557 | 0.13456 | -5.09492 | 3.49E-07 | 5.1E-05 |
| FABP5 | 113.9944 | -0.64324 | 0.130358 | -4.9344 | 8.04E-07 | 0.000107 |
| PLK1 | 94.39407 | -0.60193 | 0.123293 | -4.88212 | 1.05E-06 | 0.00013 |
| AURKA | 63.49402 | -0.7965 | 0.163934 | -4.85868 | 1.18E-06 | 0.000144 |
| XCL1 | 230.5397 | -0.87918 | 0.181734 | -4.83772 | 1.31E-06 | 0.000157 |
| CDC20 | 119.4076 | -0.58653 | 0.124917 | -4.69533 | 2.66E-06 | 0.000259 |
| RGS16 | 45.42847 | -0.80737 | 0.172204 | -4.68846 | 2.75E-06 | 0.000264 |
| HJURP | 49.45906 | -0.74368 | 0.15953 | -4.66169 | 3.14E-06 | 0.000296 |
| CCNA2 | 101.3999 | -0.60692 | 0.133681 | -4.5401 | 5.62E-06 | 0.00049 |
| SPAG5 | 74.54383 | -0.5929 | 0.135065 | -4.38979 | 1.13E-05 | 0.000881 |
| CCNB2 | 90.07022 | -0.60526 | 0.137944 | -4.38769 | 1.15E-05 | 0.000881 |
| PDLIM4 | 52.97124 | -0.79408 | 0.181356 | -4.37857 | 1.19E-05 | 0.000907 |

| | | | | | | |
|---|---|---|---|---|---|---|
| KIF23 | 60.43094 | -0.62582 | 0.143994 | -4.34614 | 1.39E-05 | 0.001038 |
| RACGAP1 | 57.86942 | -0.71329 | 0.164652 | -4.33213 | 1.48E-05 | 0.001092 |
| CDCA5 | 70.74992 | -0.593 | 0.141329 | -4.1959 | 2.72E-05 | 0.001891 |
| APOBEC3B | 60.80502 | -0.764 | 0.183197 | -4.17038 | 3.04E-05 | 0.002043 |
| KIFC1 | 30.41791 | -0.81942 | 0.203723 | -4.02224 | 5.76E-05 | 0.003506 |
| KIF20A | 47.40967 | -0.66754 | 0.167585 | -3.98331 | 6.8E-05 | 0.003894 |
| IL18R1 | 38.14404 | -0.71126 | 0.180252 | -3.94594 | 7.95E-05 | 0.004424 |
| CAV1 | 49.46489 | -0.65875 | 0.171743 | -3.83564 | 0.000125 | 0.006364 |
| IL13 | 45.44287 | -0.77858 | 0.203576 | -3.8245 | 0.000131 | 0.00649 |
| CD83 | 54.45562 | -0.63803 | 0.168937 | -3.77674 | 0.000159 | 0.007549 |
| CDC45 | 57.58178 | -0.59679 | 0.164809 | -3.62109 | 0.000293 | 0.011989 |
| HMMR | 39.17415 | -0.624 | 0.173949 | -3.58728 | 0.000334 | 0.013503 |
| LINC01281 | 42.42188 | -0.59939 | 0.168103 | -3.56559 | 0.000363 | 0.014335 |
| EXO1 | 33.29814 | -0.69232 | 0.196385 | -3.52532 | 0.000423 | 0.015948 |
| TYMS | 171.4017 | -0.68717 | 0.200636 | -3.42497 | 0.000615 | 0.020651 |
| CDCA2 | 31.56426 | -0.67213 | 0.197156 | -3.40911 | 0.000652 | 0.021519 |
| NR4A2 | 31.7803 | -0.7202 | 0.212016 | -3.39692 | 0.000681 | 0.022375 |
| TROAP | 34.27303 | -0.6042 | 0.181153 | -3.33533 | 0.000852 | 0.026205 |
| LMNB2 | 34.4321 | -0.6087 | 0.187651 | -3.2438 | 0.001179 | 0.033139 |

Supplemental Table 4: Antibody resource table

| Reagent Type | Designation | Source | Category Number | Host and subclass | Clone name |
|---|---|---|---|---|---|
| Antibody | Anti-human CD3 | BioLegend | 317330 | Mouse IgG2a, k | OKT3 |
| Antibody | Purified anti-human CD3 | BioLegend | 317302 | Mouse IgG2a, k | OKT3 |
| Antibody | Anti-human CD4 | BioLegend | 344622 | Mouse IgG1, k | SK3 |
| Antibody | Anti-human CD4 | BioLegend | 317450 | Mouse IgG2a, k | OKT4 |
| Antibody | Anti-human CD8 | BioLegend | 344732 | Mouse IgG1, k | SK1 |
| Antibody | Anti-human CD8 | BioLegend | 344712 | Mouse IgG1, k | SK1 |
| Antibody | Anti-human SIRPG | BioLegend | 336606 | Mouse IgG1, k | LSB2.20 |
| Antibody | Anti-human CD45RO | BioLegend | 304222 | Mouse IgG2a, k | UCHL1 |
| Antibody | Anti-human CD45RA | BioLegend | 304120 | Mouse IgG2a, k | HI100 |
| Antibody | Anti-human CD27 | BioLegend | 986910 | Mouse IgG1, k | M-T271 |
| Antibody | Anti-human CD28 | BioLegend | 302928 | Mouse IgG1, k | CD28.2 |
| Antibody | Ultra-LEAF™ Purified anti-human CD28 Antibody | BioLegend | 302934 | Mouse IgG1, k | CD28.2 |
| Antibody | Anti-human CD62L | BioLegend | 304809 | Mouse IgG1, k | DREG-56 |
| Antibody | Anti-human CCR7 | BioLegend | 353223 | Mouse IgG2a, k | G043H7 |
| Antibody | Anti-human GZMK | BioLegend | 370508 | Mouse IgG1, k | GM26E7 |
| Antibody | Anti-human GZMB | BioLegend | 515406 | Mouse IgG1, k | GB11 |

Supplemental Table 5: qPCR primers resource table

| Gene name | Forward primer | Reverse primer |
|---|---|---|
| UBE2C | GATGACCCTCATGGCAGTGG | TTCTCTGGGACCGGACAGTA |
| TOP2A | GGGGTCCTGCCTGTTTAGTC | AGGCTGCAATGGTGACACTT |
| CCNB1 | CCCCTGCAGAAGAAGACCTG | AGTGACTTCCCGACCCAGTA |
| CRTAM | TGTGCCTAACGTAACCCTGC | TGAAAGGAGTTGCCAGCACA |
| TNFRSF18 | CACCCAGTTCGGGTTTCTCA | ACATGCACTGACTCCTCAGC |
| XCL1 | TCTGGCTAGTGTCTATCAGAGGT | ATGGGAACCCAGTGAAGACT |
| GAPDH | GACAGTCAGCCGCATCTTCT | GCGCCCAATACGACCAAATC |
